# Atria: An Ultra-fast and Accurate Trimmer for Adapter and Quality Trimming

**DOI:** 10.1101/2021.09.07.459340

**Authors:** Jiacheng Chuan, Aiguo Zhou, Lawrence Richard Hale, Miao He, Xiang Li

## Abstract

**Background:** As Next Generation Sequencing takes a dominant role in terms of output capacity and sequence length, adapters attached to the reads and low-quality bases hinder the performance of downstream analysis directly and implicitly, such as producing false-positive single nucleotide polymorphisms (SNP), and generating fragmented assemblies. A fast trimming algorithm is in demand to remove adapters precisely, especially in read tails with relatively low quality.

**Findings:** We present a trimming program named Atria. Atria matches the adapters in paired reads and finds possible overlapped regions with a super-fast and carefully designed byte-based matching algorithm (*O(n)* time with *O(1)* space). Atria also implements multi-threading in both sequence processing and file compression and supports single-end reads.

**Conclusions:** Atria performs favorably in various trimming and runtime benchmarks of both simulated and real data with other cutting-edge trimmers. We also provide an ultra-fast and lightweight byte-based matching algorithm. The algorithm can be used in a broad range of short-sequence matching applications, such as primer search and seed scanning before alignment.

**Availability & Implementation:** The Atria executables, source code, and benchmark scripts are available at https://github.com/cihga39871/Atria under the MIT license.

## Statement of Need

### Background

Next generation sequencing (NGS) is a revolutionary new technology that produces massive, high-resolution genome sequence data to facilitate a broad range of biological applications. Illumina paired-end sequencing can read a DNA fragment from both ends and generate accurate reads for downstream bioinformatics analysis, such as assembly, resequencing, transcriptome profiling, variant calling, epigenome profiling, chromatin interaction, and chromosomal rearrangements [1, 2].

In paired-end library preparation, adapter sequences are the technical sequences ligated to both sides of inserts, which are the DNA fragments of interest. Then, DNA molecules with adapters are sequenced from both ends of the inserts so paired-end reads are generated. If insert sizes of paired-end reads are less than the read lengths, inserts are reversely complementary, and adapters are sequenced after reading through the inserts (**Fig. 1**). Thus, adapter contamination in the 3’ end needs to be removed before downstream analysis.

**Figure 1.**
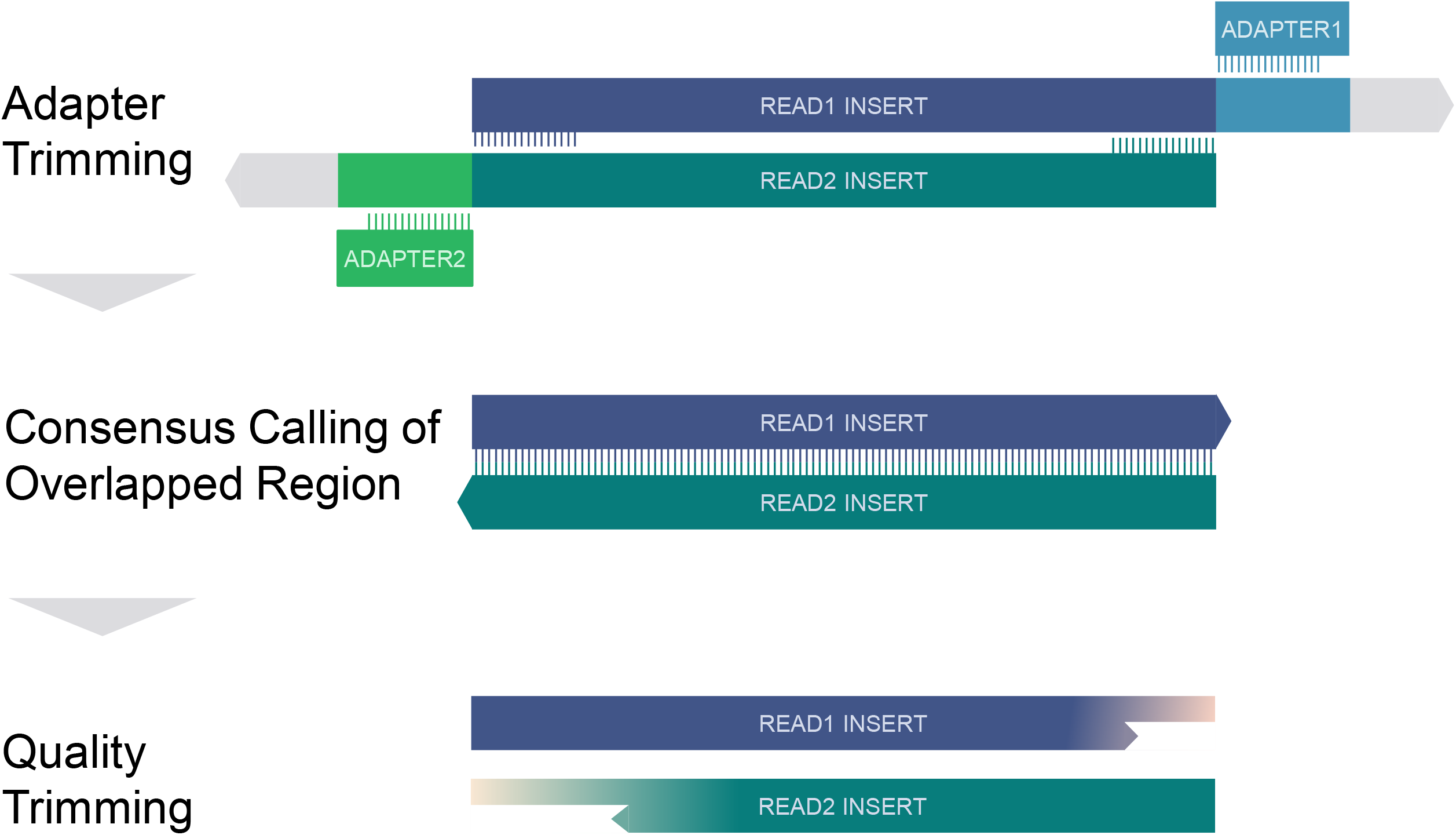
Overview of Atria workflow.

Cleaning adapters can therefore be achieved by searching adapter sequences and/or aligning paired reads (**Fig. 1**). To date, some trimmers, such as AdapterRemoval [3], Trim Galore [4], and Trimmomatic [5], use both types of information to clean adapters. However, when the quality of sequencing reads decreases, the trimming process employing both types of information is likely to give different trimming suggestions. Trimmers thus face a bottleneck when working on trimming adapters at accurate positions. Also, extremely short adapters at the low-quality 3’ end are sometimes difficult to detect. Thus, trade-offs between trimming truncated adapters, and retaining inserts intact, become necessary.

These two issues hinder trimmers from cleaning adapter sequences and leaving DNA inserts intact. To combat this, we launch Atria, an integrated trimming program for NGS data. Atria uses a super-fast byte-based matching algorithm to detect adapters and reverse complementary regions of paired reads, and integrates carefully designed decision rules to infer true adapter positions. Thus, Atria can trim extremely short adapter sequences at accurate positions and not over-trim reads without adapters (**Fig. 1**).

In addition to adapter trimming, Atria integrated a set of trimming and filtering methods, such as consensus calling for overlapped regions, quality trimming, homopolymer trimming, N trimming, hard clipping from both ends, and read complexity filtration.

### Implementation

The adapter finding algorithms used in Atria can be categorized in the following portions: DNA encoding, matching algorithm, matching and scoring, decision rules, consensus calling, quality trimming, and IO optimization (**Fig. 2**).

**Figure 2.**
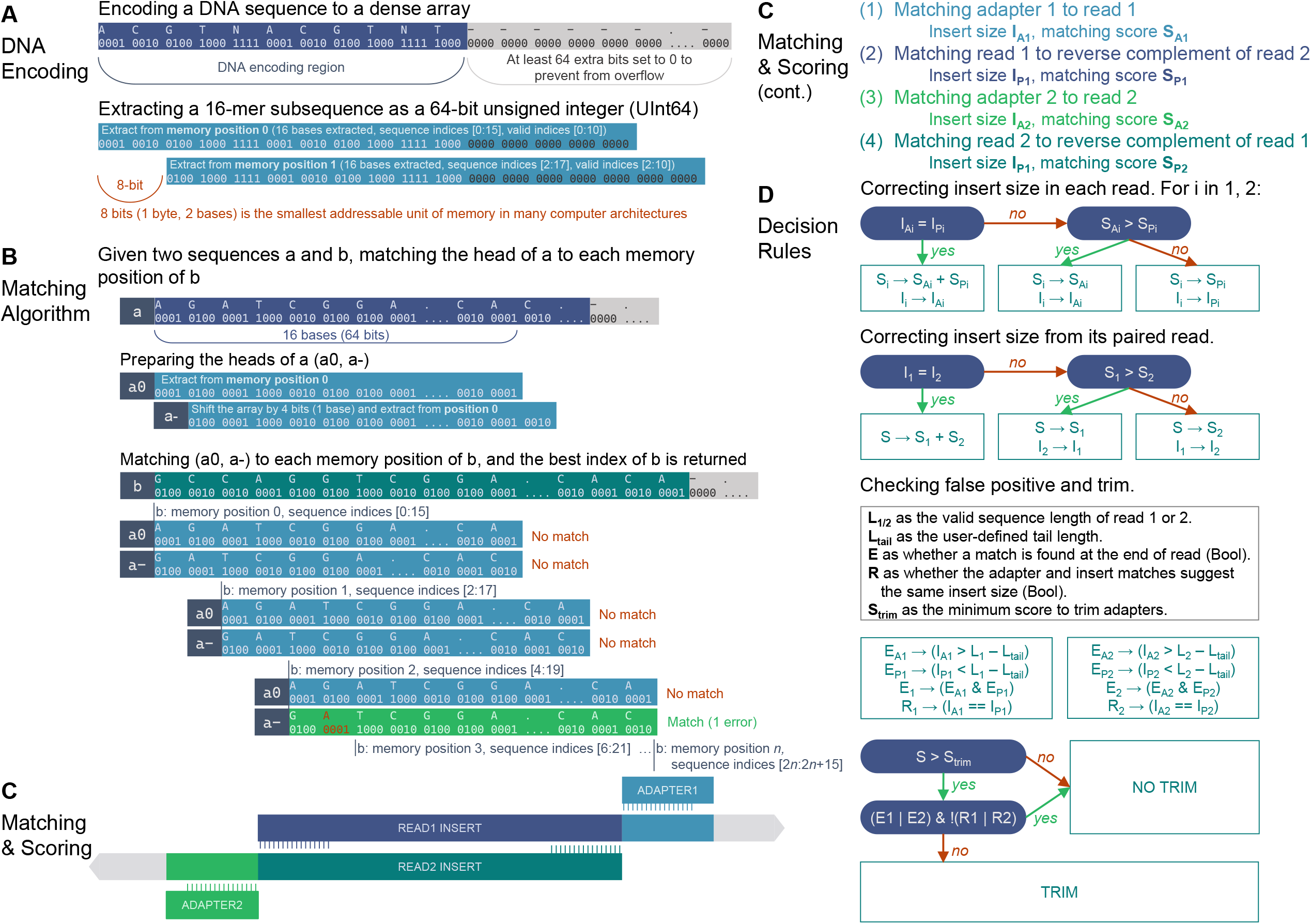
Adapter trimming algorithms.

#### DNA encoding

The DNA encoding algorithm is developed based on BioSequences, a Julia package from BioJulia [6]. The original BioSequences package encodes DNA bases A, C, G, T as four-bit codes 0001, 0010, 0100, 1000, respectively. Extended codes are also supported, such as N (1111), S (0110), and gap (0000). DNA sequences are encoded and stored in a contiguous block of Random-Access Memory (RAM) as a dense array of unsigned 64-bit integers (UInt64) (**Fig. 2A**).

Atria makes use of the property of dense arrays to extract sequences as unsigned integers from available memory locations. When accessing the last several indices of a sequence, the extraction is illegal because operating systems do not allow the loading of data outside of sequence boundary. To solve the issue, Atria constructs a bit-safe sequence array, which elongates the sequence boundary by appending a UInt64 to the end of the original array, and setting all bits after the end of encoded DNA to 0 (**Fig. 2A**).

It is noticeable that the smallest addressable unit of memory is one byte (8 bits) while each DNA is encoded in four bits, so only the even indices of sequence can be directly extracted (defining indices start from 0) (**Fig. 2A**). The extraction from odd indices requires extra operations, which is avoidable in many scenarios of a well-designed algorithm.

We denote a UInt64 extracted from the memory position *n* of sequence *a* by *a*_*n*_. *a*_*n*_ is a 16-mer and represents the subsequence of *a* indexed from *2n* to *2n*+15, which is denoted by *a*[2*n*:2*n+*15] (**Fig. 2A**).

#### Matching algorithm

Given two sequences *a* and *b*, we plan to match the 16-base-long head of *a* to each index of *b*. However, only the even indices of *b* can be extracted from memory without bitwise operations, so we prepared two UInt64 of *a*: *a*_*0*_ and *a*_*-*_. *a*_*0*_ is the 16-mer UInt64 loaded from the position 0 of *a*, and *a*_*-*_ can be computed from the following bitwise operations: (*a*_*0*_ *>> 4* | *a*_*1*_ << *4*). In this way, *a*_*0*_ represents the subsequence of *a* indexed from 0 to 15 (*a*[0:15]), and *a*_*-*_ represents *a*[1:16] (**Fig. 2B**).

In this way, the problem of matching the 16-base-long head of *a* to each index of *b* is converted to the problem of matching two 16-mers, *a*_*0*_, and *a*_*-*_, to each addressable memory position of *b*. The latter requires less bitwise operations.

The number of mismatches *K* is computed in the formula:

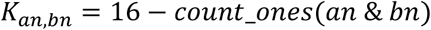

where *count_ones* counts the number of ones in the binary representation of the UInt64.

Let *k* denote the user-defined number of mismatches allowed in the 16-mer comparison of UInt64 *a*_*n*_ and *b*_*n*_ (*k =* 2 by default). After matching *a*_*0*_ and *a*_*-*_ to each addressable memory position of *b*, if the minimum number of mismatches is not greater than *k*, the smallest index of *b* of the minimum mismatches is reported.

Therefore, the complexity of the matching algorithm is *O(n)* time with *O(1)* space, so its speed is extremely fast. One limitation is that when computing the number of mismatches of *a*_*n*_ and *b*_*n*_, and if they have ambiguous bases in the same indices, the number of mismatches is underestimated. Another limitation is that the algorithm does not handle indels. Those limitations are compensated in the design of adapter matching, scoring, and decision rules.

#### Matching and scoring

We implement four pairs of matching to utilize properties of paired-end reads thoroughly: (1) matching adapter 1 head to read 1, (2) matching adapter 2 head to read 2, (3) matching read 1 head to reverse complement of read 2 and (4) matching read 2 head to reverse complement of read 1 (**Fig. 2C**). If the maximum number of bases matched of (1) and (2) is less than a user-defined cut-off (default is 9), (3) and (4) will be performed with a loosed *k* (= *k*_*original*_ *+* 1). If the largest number of matched bases of the four matches is greater than the cut-off, and some matches do not meet the requirement, we will re-run those matches with a loosed *k* (= *k*_*original*_ *+* 3) at the insert size indicated from the best match. If the new number of matched bases is greater than the cut-off, the old match will be discarded.

The scoring system measures the matching reliability of the whole 16-mer rather than each base. The Phred quality score *Q* of each base is converted to the probability *P* of that the corresponding base being correct using the formula:

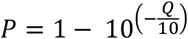

Then, the average base quality 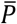 of 16-mer sub-sequence *a* at the memory position *n* is computed:

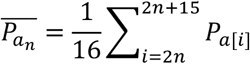

Notably, if the read quality is too low, it would imply an invalid match. However, in reality, invalid matches are filtered out by the kmer-based algorithm. To solve the discordance, we limit the lower bound of 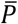 to 0.75 manually.

The matching score *S* between *a*_*n*_ and *b*_*m*_ is defined as:

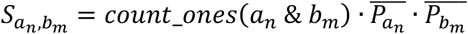

where *count_ones* counts the number of ones in the binary representation of the UInt64. When sequence *a* is a user-defined adapter, 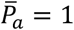 is used. Generally, the matching score *S* is ranged from 0 to 16.

### Pseudocode 1: Matching and scoring

~~~
r1_pos_adpt, r1_nmatch_adpt = match(adapter1, r1, k)
r2_pos_adpt, r2_nmatch_adpt = match(adapter1, r1, k)
k_extra = max(r1_nmatch_adpt, r2_nmatch_adpt) < 9 ? 1 : 0
r1_pos_pe, r1_nmatch_pe = match(reverse_complement(r2), r1, k + k_extra)
r2_pos_pe, r2_nmatch_pe = match(reverse_complement(r1), r2, k + k_extra)
max_nmatch = max(r1_nmatch_adpt, r2_nmatch_adpt, r1_nmatch_pe, r2_nmatch_pe)
max_pos = corresponding position of max_nmatch
if max_nmatch > 9
  for matches with any nmatch < 9
    redo match with loosed k = k + 3 at max_pos
    replace old results if nmatch > 9
r1_prob_adpt = average_16mer_quality(r1, r1_pos_adpt)
r2_prob_adpt = average_16mer_quality(r2, r2_pos_adpt)
r1_prob_head = average_16mer_quality(r1, 1)
r2_prob_head = average_16mer_quality(r2, 1)
r1_prob_pe = average_16mer_quality(r1, r1_pos_pe)
r2_prob_pe = average_16mer_quality(r2, r2_pos_pe)
r*_prob_* = 0.75 if any r*_prob_* < 0.75
r1_score_adpt = r1_nmatch_adpt * r1_prob_adpt
r2_score_adpt = r2_nmatch_adpt * r2_prob_adpt
r1_score_pe = r1_nmatch_pe * r1_prob_pe * r2_prob_head
r2_score_pe = r2_nmatch_pe * r2_prob_pe * r1_prob_head
~~~

#### Decision rules

This module infers correct adapter positions from the four pairs of matching described in the previous section. It is illustrated and self-explanatory in **Fig. 2D**. First, in each read, the adapter and paired-end matches are compared. The one with the higher matching score is chosen. If both matches support the same adapter position, the matching score of the read is the sum of adapter and paired-end matching scores. Then, the matches of the two paired-end reads are compared using the same strategy. If one read finds an ideal adapter (matching score > 10 by default) while the other read is too short to check or the average base accuracy of its 16-mer is less than 0.6 (Phred Q < 5), both reads will be trimmed. If the matching score of a given read pair is less than 10 (by default), the read pair will not be trimmed.

Other read pairs will be taken a further examination to reduce false positives, which are usually adapter matches at read tails. A read tail is defined as the last several bases (default is 12 bp) of each read. Reads are not trimmed if both statements are true: (1) In any paired read, the adapter is found at the tail, but the paired-end match is not; (2) In both paired reads, adapter and pair-end matches suggest different trimming positions.

Before the final trimming, one additional step is required for the accurate positioning of adapter sequences. The previous steps usually assume the read 1 and 2 have the same length of insert sizes, but indel in reads usually lead to over or under trim one base. To prevent this circumstance, Atria re-positions the adapter by matching one adjacent base with the first four bp of adapter sequences. The position of the highest number of bases matched is chosen to trim. This step is ignored when the inferred insert size is greater than the read length minus three because, in this situation, the adapter sequence is too short to check.

### Pseudocode 2: Decision rules

~~~
function correct_insert_size(pos1, score1, pos2, score2)
  if pos1 == pos2
    return pos1, score1 + score2
  else
    score = max(score1, score2)
    pos = corresponding pos of max score
    return pos, score
r1_pos, r1_score = correct_insert_size(r1_pos_adpt, r1_score_adpt, r1_pos_pe, r1_score_pe)
r2_pos, r2_score = correct_insert_size(r2_pos_adpt, r2_score_adpt, r2_pos_pe, r2_score_pe)
r12_pos, r12_score = correct_insert_size(r1_pos, r1_score, r2_pos, r2_score)
if r1_pos != r2_pos
  if r1_score > 10
    r2_prob = average_16mer_quality(r2, r1_pos)
    @goto “trim” if r2_prob < 0.6
  elseif r2_score > 10
    r1_prob = average_16mer_quality(r1, r2_pos)
    @goto “trim” if r1_prob < 0.6
function check_read_tail(read)
  E_adpt = whether adapter found at read tail
  E_pe = whether pair-end match found at read tail
  E = E_adpt & E_pe # both matches in read tail
  R = rx_pos_adpt == rx_pos_pe # adapter and pair-end match at same position return E, R
E1, R1 = check_read_tail(r1)
E2, R2 = check_read_tail(r2)
E = E1 | E2 # at least one read matching in read tail
R = R1 | R2 # at least one read matching at the same position
is_false_positive = E & !R
if r12_score > trim_score & !is_false_positive
  @label “trim”
  r1_pos_adjusted = adjacent_one_bp_check(r1, adapter1, r12_pos)
  r2_pos_adjusted = adjacent_one_bp_check(r2, adapter2, r12_pos)
  trim(r1, r1_pos_adjusted)
  trim(r2, r2_pos_adjusted)
~~~

#### Consensus calling

In this module, the overlapped base pairs of read 1 and 2 are corrected to the corresponding bases with higher quality scores. It has three steps, prediction, assessment, and correction.

In the prediction step, Atria makes a preliminary estimate of whether a read pair contains an overlapped region. If adapters are trimmed and the remaining lengths of read 1 and 2 are the same, the prediction passes. If no adapter can be trimmed, two additional matching and scoring are required. The head of the reverse complement of read 2 is matched to read 1, and the head of the reverse complement of read 1 is matched to read 2. If the two matches reach a consensus, the prediction passes. Otherwise, the prediction fails and consensus calling is skipped.

In the assessment step, Atria compares the whole overlapped region using a similar matching algorithm, except that ambiguous bases (N, 1111) are converted to gaps (0000) before matching. If the ratio of mismatch is greater than a user-defined value (28% by default), the assessment fails, and consensus calling is skipped.

In the correction step, each base pair in the overlapped region is corrected to the corresponding base with the highest quality score.

#### Quality trimming

Atria implements a traditional sliding window algorithm to remove the low-quality tail. The sliding window scans from the front of the read and computes the average Phred quality score of the sliding window. If the average quality is less than a given threshold, the read tail is removed.

#### IO optimization

Atria spends more time on reading and writing than matching and trimming, so the key to reducing runtime is to optimize IO usage. Considering that a large amount of RAM is easily accessible nowadays, Atria trades increased RAM usage with decreased time. A large block of memory is allocated for reading input files, which is then wrapped and encoded to FASTQ objects parallelly using multi-threading. On the contrary, in the writing process, Atria unboxes and decodes FASTQ objects to string vectors in parallel and writes sequentially to files. In addition, pigz (parallel gzip) and pbzip2 (parallel bzip2) are called for compression and decompression when needed [7, 8]. Atria also support running with a single thread.

## Comparison to related work

### The performance of adapter trimming on a simulated dataset

We simulated 8.9 G bases with 100 bp paired-end reads from the *Arabidopsis thaliana* reference genome using the Skewer modified ART, a public NGS read simulator to allow adapters in the reads [9, 10]. The simulation profile was trained from a 101 bp paired-end public dataset SRR330569, and the 33 bp adapter pair used in read simulation is AGATCGGAAGAGCACACGTCTGAACTCCAGTCA and AGATCGGAAGAGCGTCGTGTAGGGAAAGAGTGT [11].

Atria v3.0.0 was benchmarked with cutting-edge and popular trimmers, including AdapterRemoval v2.3.1 [3], Skewer v0.2.2 [10], Fastp v0.21.0 [12], Ktrim v1.2.1 [13], Atropos v1.1.29 [14], SeqPurge v2012_12 [15], Trim Galore v0.6.5 [4] and Trimmomatic v0.39 [5]. Only adapter trimming was used, and other trimming and filtration were disabled. Detailed command line arguments are listed in **Table S1**. Each trimming software was running on an idle Ubuntu 19.10 server with a 32-thread Intel i9-9960X Central Processing Unit (CPU) @ 3.10 GHz, 128 gigabyte (GB) DDR4-3200 RAM, and a 2 terabyte (TB) Samsung 970 EVO Solid State Drive (SSD) (sequential reads and writes up to 3.5 and 2.5 TB/s).

The trimming performance was evaluated based on the following metrics: positive predictive value (PPV), as the fraction of the number of correctly trimmed reads to all trimmed reads; sensitivity, as the fraction of the number of correctly trimmed reads to the reads with adapters; specificity, as the fraction of the number of untrimmed reads without adapters to all reads without adapters; and Matthew’s correlation coefficient (MCC) measuring overall quality of pattern recognition, as

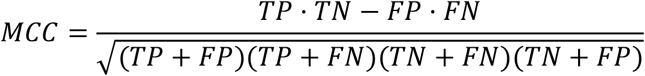

where TP is the number of reads trimmed correctly, TN is the number of untrimmed reads without adapters, FP is the number of over-trimmed reads, and FN is the number of under-trimmed reads [3, 10].

The adapter trimming performance is shown in **Table 1**. AdapterRemoval, Atria and Skewer were the top-class adapter trimmers in terms of MCC (99.61%, 99.51%, 99.44%, respectively) (**Table 1**). Fastp (98.92%) and Atropos (98.00%) were in the second tier (**Table 1**). Ktrim obtained a good specificity (98.84%) but sacrificed its sensitivity (85.85%), and Trimmomatic achieved an exceptional specificity (99.88%) by trading off its sensitivity (57.86%) (**Table 1**).

**Table 1.**
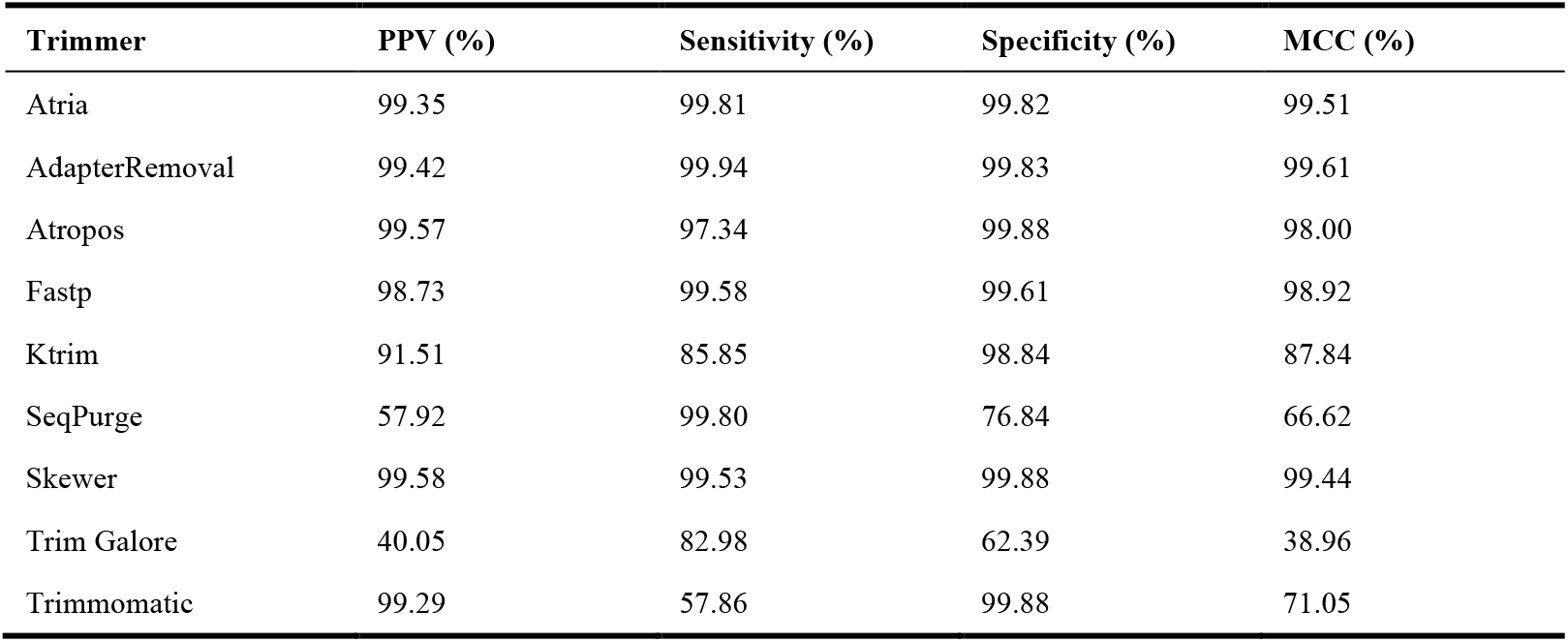
Adapter trimming performance on the 8.9 G bases with 100 bp paired-end simulated data.

To compare speed and efficiency, elapsed time (wall time) and average CPU consumption of each trimmer were recorded in different threading (1-32 threads) for uncompressed and gzip compressed data formats (**Fig. 3, Table S1**). Efficiency was defined as the fraction of processing speed to the percent of CPU utilized, so it was a better measurement, especially in CPU-intensive scenarios, such as running on a server with a job scheduling system or trimming multiple samples at the same time. Ktrim and Atria were two of the fastest trimmers in terms of speed and efficiency, from one to 16 threads (**Fig. 3, Table S1**). For uncompressed data, Trimmomatic was faster than Atria using 8-32 threads, but its real CPU usage was much greater than Atria (**Fig. 3, Table S1**). The speed and efficiency of AdapterRemoval and Skewer were generally 2-4 times less than Atria, and Atropos was the slowest one (**Fig. 3, Table S1**). SeqPurge did not support the output of uncompressed data, so it was only tested in the compressed benchmark.

**Figure 3.**
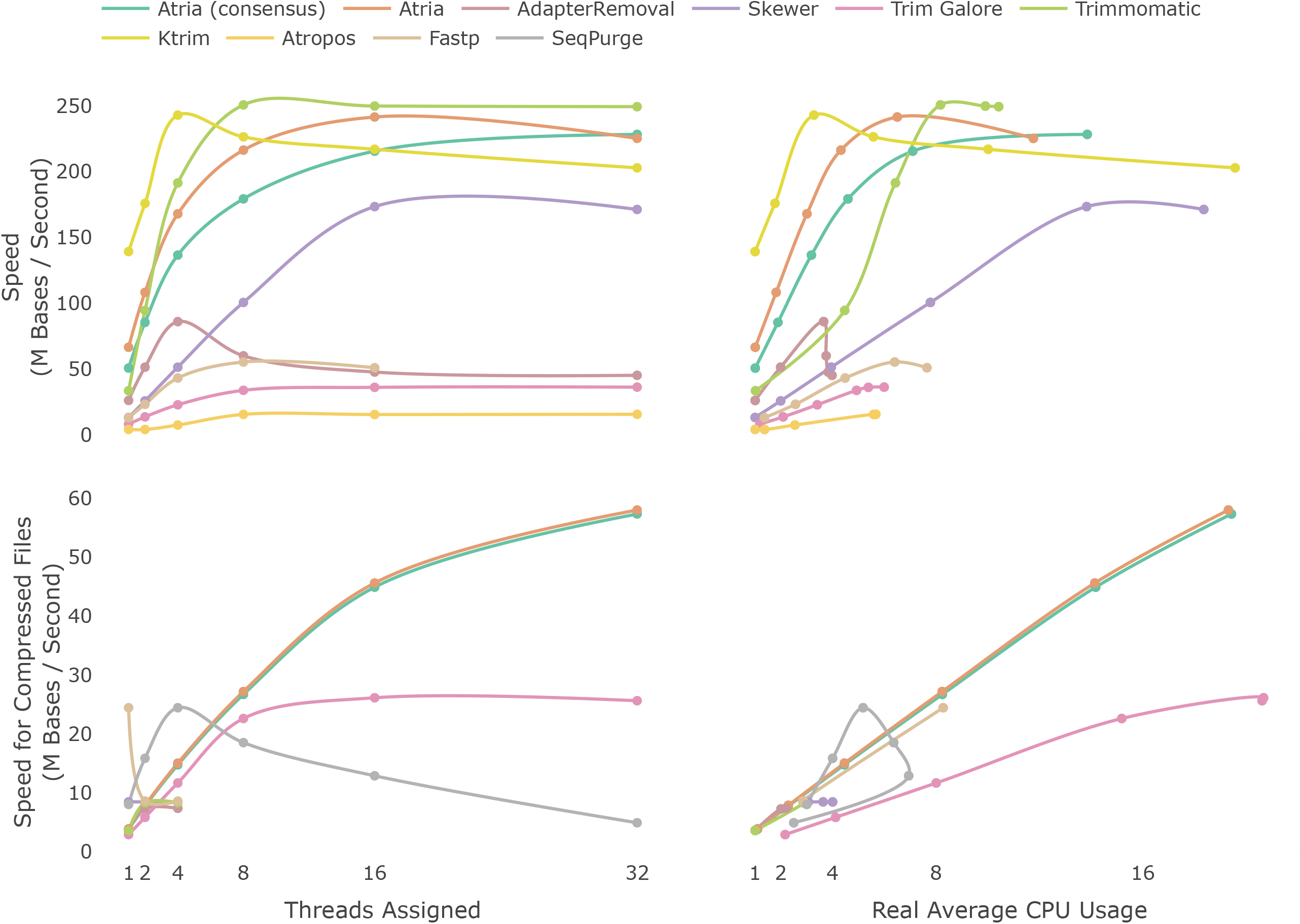
Benchmark of adapter-trimming speed for uncompressed and compressed files on different threading options. The 8.9 G bases simulated paired-end data with a 100 bp read length was trimmed in both uncompressed and compressed format using up to 32 threads. Speed is the ratio of the number of bases to elapsed time (wall time). SeqPurge does not support uncompressed outputs, so it is not shown in the uncompressed benchmark. In the trimming for compressed data, the speed of AdapterRemoval, Skewer, Fastp, Atropos, and Trimmomatic kept constant when the number of threads increased from 4 to 32, so we only benchmark those trimmers using 1, 2, and 4 threads. Ktrim does not support output compressed files, so it is not shown in the compressed benchmark.

When trimming compressed data, the speed of AdapterRemoval, Skewer, Fastp, Atropos and Trimmomatic kept constant when the number of threads increased from 4 to 32, because they failed to utilize more than four CPU in the IO process, while Atria and Trim Galore did not have the limitation (**Fig. 3, Table S1**). Atria was faster than Trim Galore, and the efficiency of Atria was constantly two to three times greater than Trim Galore (**Fig. 3, Table S1**). SeqPurge showed strange speed curves; when assigning a single thread to SeqPurge, the average CPU usage was 300%, and the speed and average CPU usage dropped when assigning 8 to 32 threads (**Fig. 3, Table S1**). In addition, Ktrim did not support output compressed files, so we ignored it. In general, Atria was the fastest trimmer when trimming compressed files.

### The detailed statistics of adapter trimming accuracy on a simulated dataset

The previous portion benchmarks on a whole dataset. This section evaluates trimming accuracy regarding different read properties, including adapter presence or absence, base error, and adapter length. To achieve the goal, Atria integrates a benchmarking toolkit for read simulation and trimming analysis.

The read simulation method was inspired by how sequencers read DNA. First, an original DNA fragment (insert) with a given original insert size is simulated base by base. Adenine, thymine, cytosine, and guanine are randomly chosen repetitively. Then, the insert and adapter sequences are copied base by base with an error profile, which simulates the procedure of sequencing by synthesis. The error profile defines substitution rate, insertion rate, and deletion rate.

Twenty-one million read pairs were simulated with a uniform read length (100 bp), different error profiles, adapter length, and original insert sizes. The baseline error profile comprises a 0.1% substitution rate, 0.001% insertion rate, and 0.001% deletion rate, inspired by an Illumina error profile analysis [16]. 1x, 2x, 3x, 4x, and 5x baseline error profile, 16, 20, 24, 28, and 33 adapter lengths, and 66 to 120 even insert sizes are chosen. In this way, the reads with the least insert size have full lengths of adapters. The reads with 66-98 original insert sizes contain adapters, and the reads with 100-120 original insert sizes are free from adapter contamination, except for few reads with a 100 bp insert size containing indels. Therefore, in each condition combination, 30 thousand read pairs were simulated to avoid random errors. The reads were trimmed with the same method described in the last section.

The average trimming performance among different conditions is shown in **Fig. 4 A**. When adapters are present, Atria trims 99.9% adapters accurately, and SeqPurge, Fastp, and Atropos follow closely with an accuracy of 99.7% (**Fig. 4 A1**). When adapters are absent, AdapterRemoval, Skewer, Trimmomatic, Atropos, and Atria successfully leave 100.0% reads intact, and Fastp falls behind with 99.8% accuracy (**Fig. 4 A2**).

**Figure 4.**
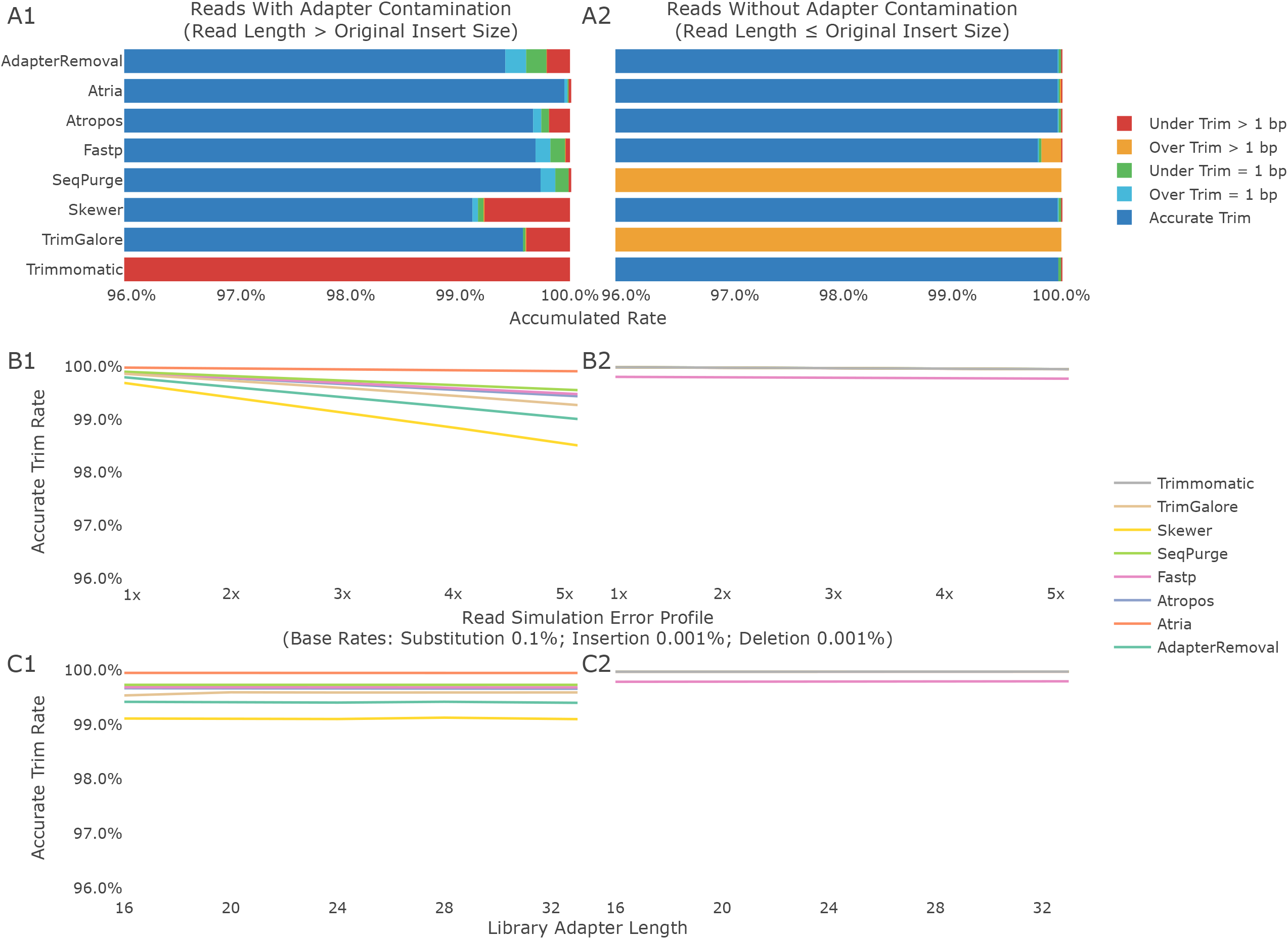
Adapter trimming accuracy on adapter presence and absence, different base errors, and adapter lengths. A1, B1, and C1 are statistics for reads with adapter contamination, while A2, B2, C2 for reads without adapters. A1 and A2 show the accumulated rates of accurate trim, one bp over trim, one bp under trim, multiple bp over trim, and multiple bp under trim. In A1, the accuracy of Trimmomatic is 41.0%. In A2, the accuracy of SeqPurge is 78.8%, the accuracy of Trim Galore is 68.3%. B1 and B2 show the trimming accuracy on different error profiles. In B1, the accuracy of Trimmomatic drops from 41.9% to 40.1%. In B2, the accuracy of SeqPurge is 78.8%, and the accuracy of Trim Galore is 68.2 - 68.3%. C1 and C2 show the trimming accuracy on different adapter lengths. In C1, the accuracy of Trimmomatic is 0.0% at 16 bp adapter length, 50.7% to 51.6% at adapter lengths from 20 to 33 bp. In C2, the accuracy of SeqPurge ranges from 78.7% at 16 bp to 78.9% at 33 bp, and the accuracy of Trim Galore ranges in 68.2 - 68.3% from 16 to 33 bp.

**Fig. 4 B** illustrates the trimming accuracy on different read error profiles. When adapters are present, the accuracy of all trimmers drops as error rates increase (**Fig. 4 B1**). Atria keeps the highest accuracy from 100.0% to 99.9%, and is almost not affected by different error rates (**Fig. 4 B1**). The accuracy of SeqPurge, Fastp, and Atropos decrease from 99.9% to 99.6%, 99.5%, and 99.4%, respectively (**Fig. 4 B1**). With no adapter present in reads, the accuracy is hardly influenced by error profiles (**Fig. 4 B2**), so the performance is similar to **Fig. 4 A2**. In addition, adapter lengths ranging from 16 to 33 bp are not relevant to most trimmers’ accuracy, including Atria (**Fig. 4 C**).

### The performance of adapter trimming on real sequencing dataset

#### RNA-Seq paired-end dataset (SRR330569)

SRR330569 is a real RNA-Seq dataset sequenced from *Drosophila simulans* with 5.46 G bases and 2 × 101 bp read length. It contains 38 bp adapter sequences AGATCGGAAGAGCGGTTCAGCAGGAATGCCGAGACCG and AGATCGGAAGAGCGTCGTGTAGGGAAAGAGTGTAGAT in read 1 and read 2, respectively. Adapter trimming was performed by different trimmers without other trimming or filtering methods. Then, a sliding-window based quality trimming was performed to remove low-quality tails (sliding window size = 5 and average Q score ≥ 15). The adapter-trimmed reads and adapter-and-quality-trimmed reads were mapped to the *Drosophila simulans* genome version 2.02 from FlyBase using Hisat2 v2.2.1, respectively [17, 18]. Mapping statistics were collected using SAMTools Stat v1.10 [19]. Skewer did consensus calling after adapter trimming, and no option was provided to disable it. To achieve benchmark parity, Skewer was compared to Atria with consensus calling enabled, and other trimmers were compared to Atria without consensus calling. Time trimming was recorded in accordance with a common scenario: inputs were gzip-compressed and trimmed with eight threads, and outputs were also gzip-compressed to reduce massive disk use. All tested trimmers worked in the scenario except that Ktrim could not output gzip files **(Table 2)**.

**Table 2.**
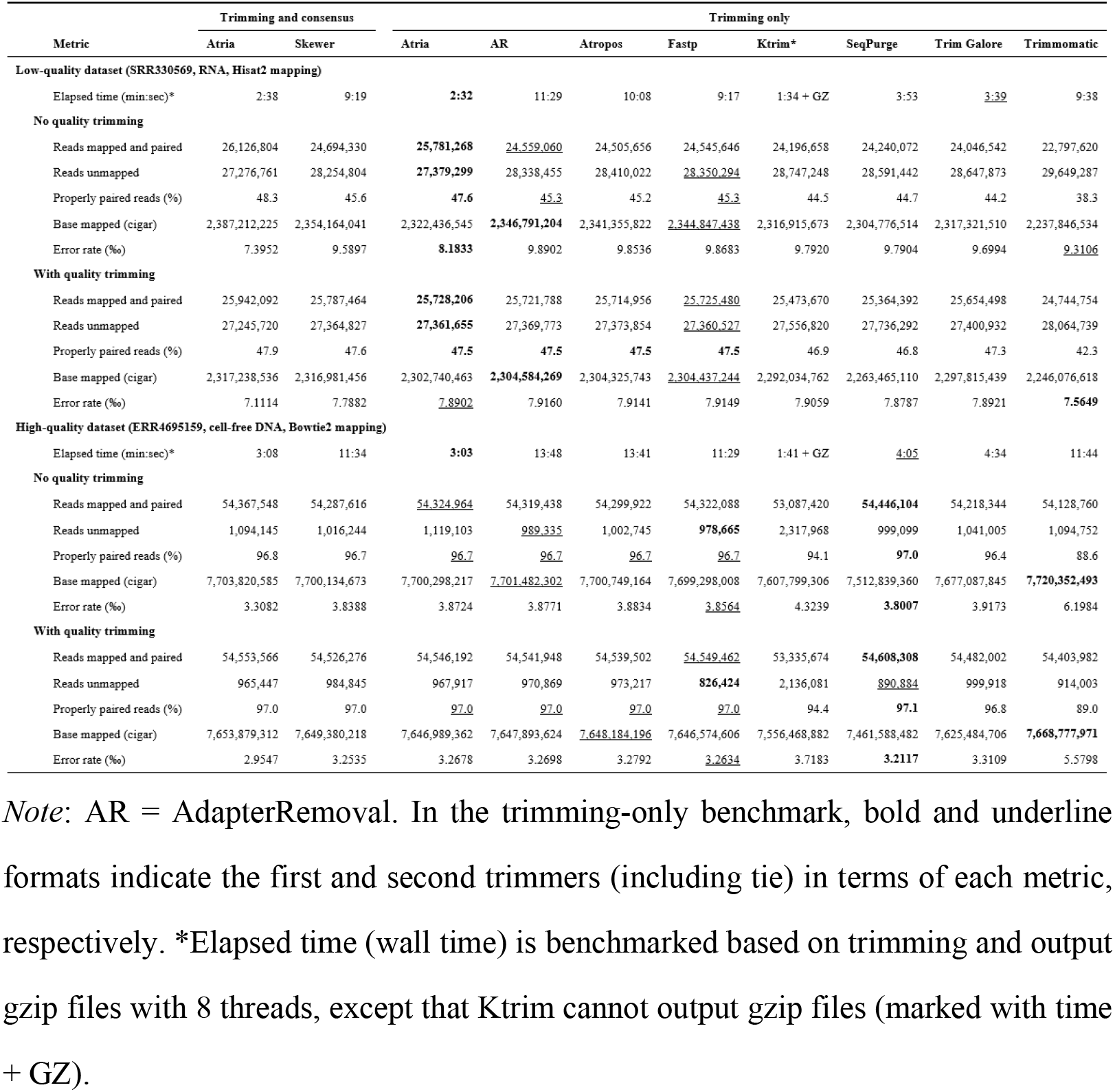
Performance of trimmers on real data.

Atria was the fastest program to process and output compressed data in terms of wall time (**Table 2**). It also achieved the highest number of reads mapped and paired, and the percent of properly paired reads with or without quality trimming. Generally, higher base mapped is accompanied with higher error rate in the mapping process, so it is important to interpret the two metrics together. Atria had the lowest mapping error rate of 8.1833‰ and the forth highest base mapped (**Table 2**). The trimmers (AdapterRemoval, Fastp, and Atropos) with the highest three error rates has the highest base mapped (**Table 2**). Our program generally improved more than 5% compared to other trimmers for the data without quality trimming (**Table 2**). The mapping statistics of data without quality trimming were generally worse than with quality trimming except for Atria. Specifically, the properly paired rates of other trimmers without quality trimming were 0.5 to 4% less than with quality trimming (**Table 2**). Quality trimming also increased the number of mapped and paired reads and reduced the number of unmapped reads (**Table 2**).

#### Genome-wide human cell-free DNA dataset (ERR4695159)

Generally, plasma cell-free DNA is short in length [20], and trimming is extremely important in medical diagnosis. Here, we chose a human genome-wide cell-free DNA dataset ERR4695159. It has 8.4 G bases with 2 × 150 bp read length with 33 bp adapter sequences AGATCGGAAGAGCACACGTCTGAACTCCAGTCA in read 1 and AGATCGGAAGAGCGTCGTGTAGGGAAAGAGTGT in read 2. The benchmark workflow was the same as the RNA-Seq analysis, except that the clean reads were mapped to the human reference genome hg38 (GRCh38.p13) using Bowtie2 v2.3.5.1 [21].

Atria was also the fastest trimmer in the scenario (3 min 3 s) (**Table 2**). SeqPurge and Trim Galore finished the task in more than 4 minutes, while others spent more than 11 minutes (**Table 2**).

In adapter-trimming-only statistics, SeqPurge had the highest mapped and paired reads (54,446,104) and the highest properly paired reads (97.0%) (**Table 2**). Atria followed with 54,324,964 mapped and paired reads. Atria, AdapterRemoval, Atropos, and Fastp all had 96.7% properly paired reads (**Table 2**).

With quality trimming, the overall performance increased, and properly paired reads were closer; SeqPurge had 97.1% properly paired reads, with Atria, AdapterRemoval, Atropos, and Fastp close behind at 97.0% (**Table 2**). Only 89.0% of reads were properly paired with Trimmomatic (**Table 2**).

## Discussion

Atria performs favorably with other cutting-edge adapter trimmers in accuracy, robustness, speed, and efficiency. Its performance is ascribed to the byte-based matching algorithm. The design concept of the algorithm is to minimize any unnecessary CPU operations by taking advantage of the data structure of dense arrays.

Matrix-based algorithms, such as the Needleman-Wunsch algorithm and the Smith-Waterman algorithm, allocate and update a matrix and perform base-to-base comparison [22, 23]. They report every mismatch and gap between two sequences while Atria skips this step since it is focussed on the start positions of successful matches. Despite that, the matching algorithm used in Atria is able to identify mismatch loci when needed.

The byte-based matching algorithm is lightweight and designed for short sequence scanning. Each DNA is encoded in four bits and stores continuously in RAM. A sub-sequence can be extracted as an unsigned integer from a given memory position. For example, a 64-bit unsigned integer (UInt) represents a 16-mer, and a 128-bit UInt represents a 32-mer. The comparison between two sub-sequences is completed within the accumulator register, a CPU unit for arithmetic or logical operation. It does not require addressing or updating a scoring matrix from RAM. When comparing a short sequence, such as an adapter, to a long sequence, such as the read, the 16-mer of the short sequence is compared to every position of the long sequence. Hence, the byte-based matching algorithm has *O(n)* expected time complexity and *O(1)* space complexity in adapter matching, where n is the length of the long sequence.

The algorithm also has its limitations. It only reports the number of matched bases and does not report the positions of mismatches, so it cannot be used for sequence alignment solely. Besides, the algorithm does not handle insert and deletion. However, the average indel rate of Illumina library is 10^−6^ to 10^−5^ [16], and the low indel rate is almost negligible in real data analysis. In addition, Atria does four pairs of matches in different locations to compensate for the limitation. If one match is failed because of indel, other matches will suggest the real adapter positions.

In the runtime benchmark, we compared how trimmers performed using extremely high CPU cores. In general, efficiency marginally decreased as CPU usage increased due to the trimmers’ parallel implementation and the inevitable cost of multi-threading, such as task scheduling and context switching. In addition, IO could be the main bottleneck for most hard disk drives and some solid-state drives. Thus, if the system IO reaches a bottleneck, an efficiency plateau would be expected sooner.

## Conclusions

We introduce not only Atria, a cutting-edge trimming software for sequence data, but also the ultra-fast and lightweight byte-based matching algorithm. The algorithm can be used in a broad range of short-sequence matching applications, such as primer search and seed scanning before alignment. Atria is implemented in Julia, a programming language designed specifically for high performance. The source code, executables, and benchmark scripts are available on Atria’s Github page [24].

## Availability and requirements

Project name: Atria

Project home page: https://github.com/cihga39871/Atria

Operating system(s): Linux, OSX

Programming language: Julia

Other requirements: Julia v1.4, Pigz v2.4 or higher, Pbzip v1.1.13 or higher

License: MIT

Research Resource Identification Initiative ID: SCR_021313

## Data Availability

The datasets SRR330569, and ERR4695159 analyzed during the current study are available in the Sequence Read Archive from the National Center for Biotechnology Information [11, 25].

The Atria source codes, releases, documents, and benchmark scripts can be downloaded from Atria’s Github page [24].

## Abbreviations

CPU: Central processing unit
DNA: Deoxyribonucleic acid
GB: Gigabyte
MCC: Matthew’s correlation coefficient
NGS: Next-generation sequencing
PPV: Positive predictive value
RAM: Random-access memory
RNA: Ribonucleic acid
SNP: Single nucleotide polymorphism
SSD: Solid-state drive
TB: Terabyte
UInt: Unsigned integer
UInt64: Unsigned 64-bit integer
WGS: Whole-genome sequencing.

## Authors’ contributions

JC developed Atria software, performed benchmark experiments under the supervision of XL. Both XL and LH serve as co-supervisors and participates in the design of the study. MH contributes to the optimization of the algorithm. AZ participates in benchmark validation. JC, LH, and XL drafted the manuscript. All authors read and approved the final version of the manuscript.

## Acknowledgments

The technical assistance of Jingbai Nie is greatly acknowledged. The financial support of CFIA to JC is greatly appreciated. The advice on algorithm optimization, encouragement, and support of Drs. Christian Lacroix and Stevan Springer to JC are greatly appreciated.

## Supplementary material

**Table S1 Trimming speed on the 8**.**9 G bases 100 bp paired-end simulated data** Atria (consensus) does both adapter trimming and paired-end consensus call (base correction of overlapped regions). In the trimming for uncompressed data, SeqPurge does not support uncompressed outputs, so it is not shown in the uncompressed benchmark. Fastp does not support 32 threads, so only 1-16 threads were tested. In the trimming for compressed data, the speed of AdapterRemoval, Skewer, Fastp, and Trimmomatic kept constant when the number of threads increased from 4 to 32, so we only benchmarked those trimmers using 1, 2, and 4 threads. Atropos was too slow to trim compressed data, and Ktrim did not support compressed outputs, so they are not shown in the compressed benchmark.

**Table 2.**
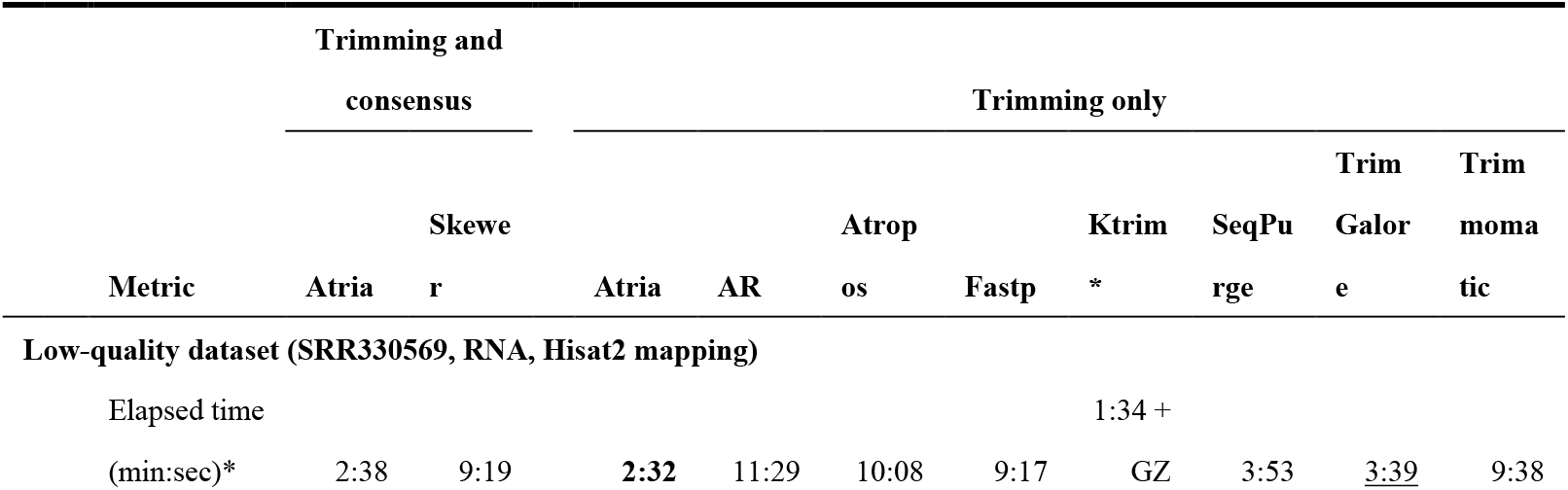
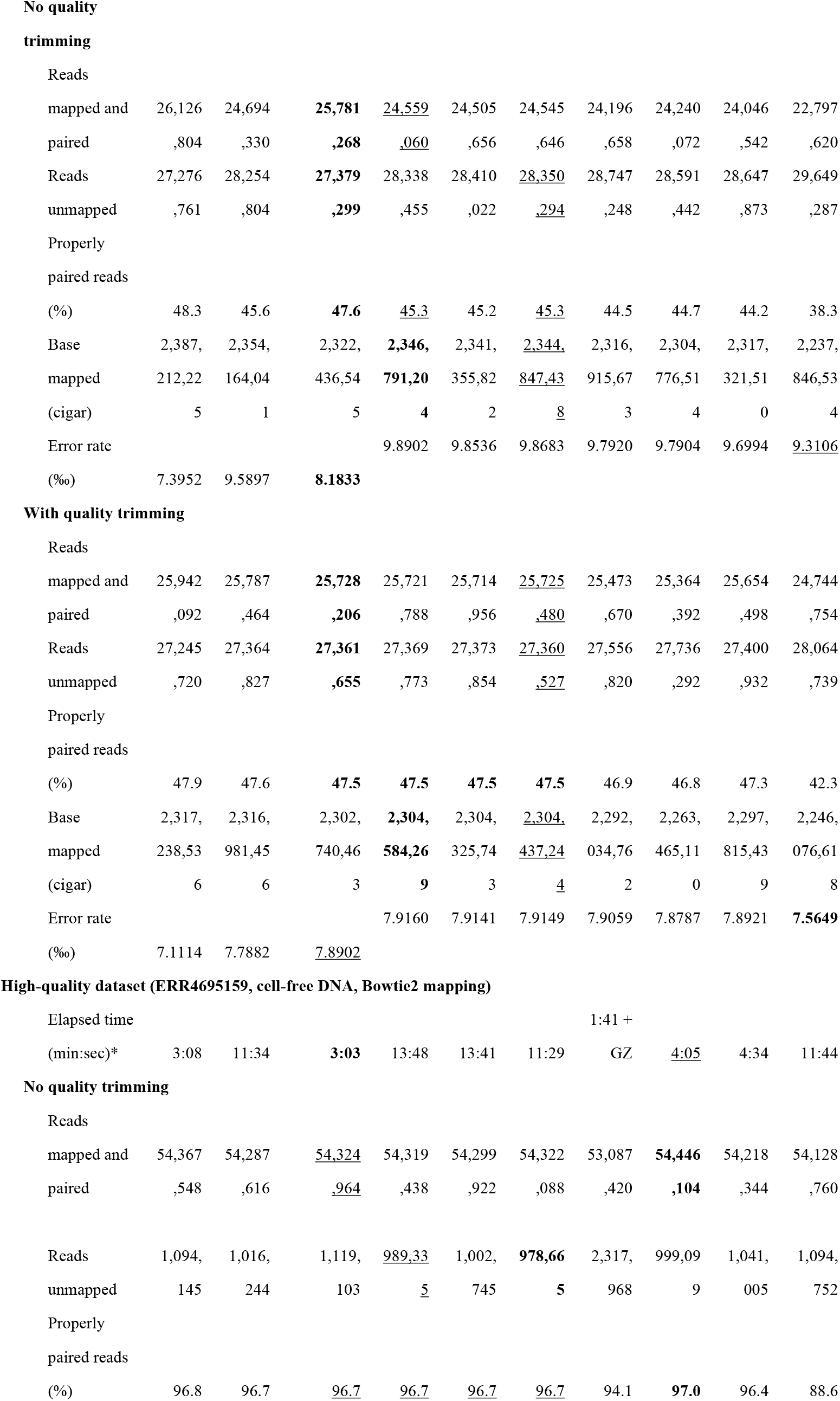
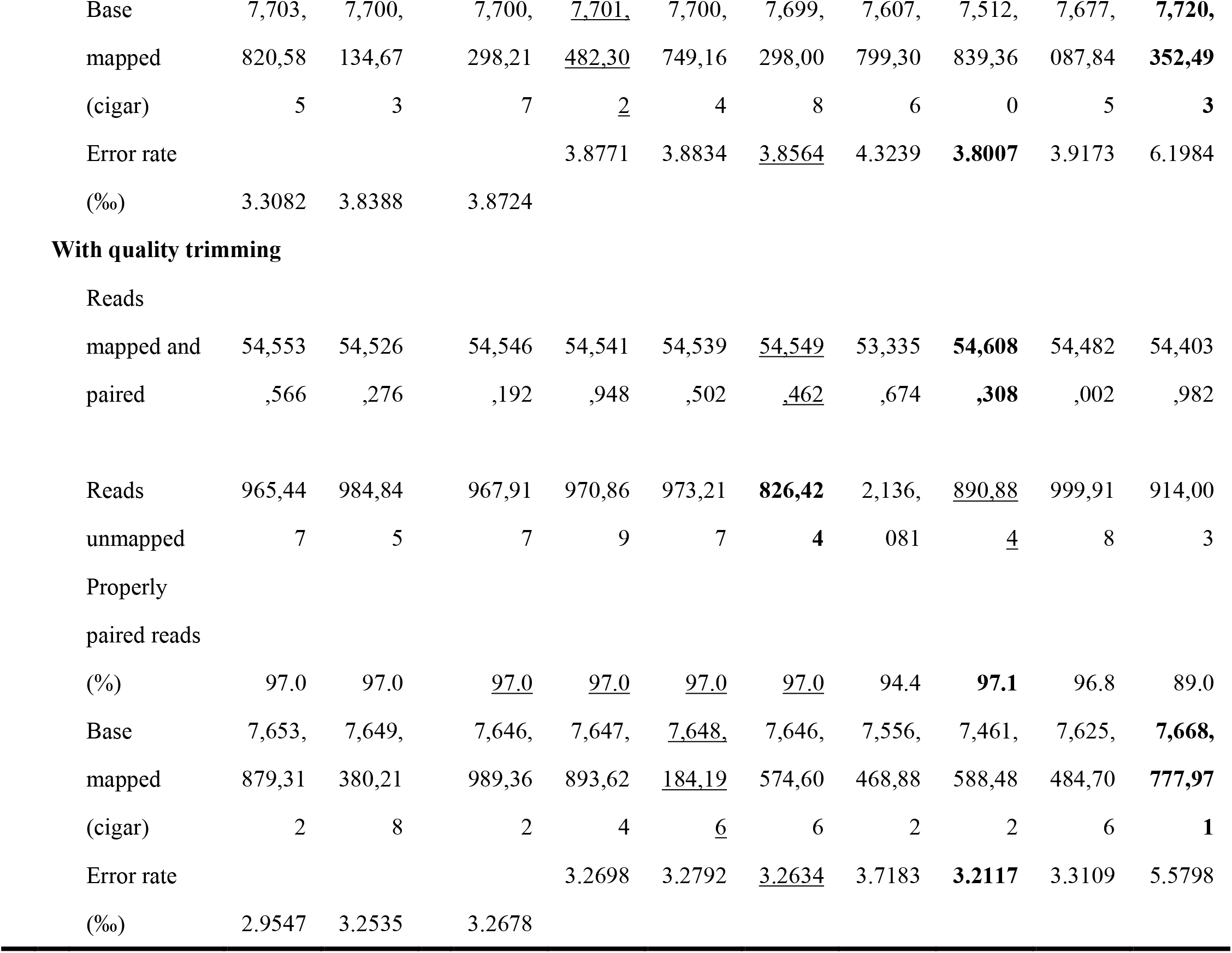
Performance of trimmers on real data (larger than A4)

