## Supplementary material for "Atria: An Ultra-fast and Accurate Trimmer for Adapter and Quality Trimming": Table S1

**Table S1-1. Trimming speed on the simulated data (uncompressed data)**

| Threads | Trimmer | Efficiency (M Bases/s/CPU) | Speed (M Bases/s) | User Time (s) | System Time (s) | CPU | Elapsed Time (s) | Max Memory (kB) | Command |
|---|---|---|---|---|---|---|---|---|---|
| 1 | Atria | 66.51 | 66.79 | 121.17 | 12.75 | 100.4% | 133.36 | 2,248,848 | atria --no-consensus -r TAIR10.sim_1.fq -R TAIR10.sim_2.fq -o Atria --no-tail-n-trim --max-n=-1 --no-quality-trim --no-length-filtration --adapter1 AGATCGGAAGAGCACACGTCTGAACTCCAGTCA --adapter2 AGATCGGAAGAGCGTCGTGTAGGGAAAGAGTGT --threads 1 |
| 1 | Atria (consensus) | 50.79 | 50.95 | 162.96 | 12.42 | 100.3% | 174.81 | 2,251,148 | atria -r TAIR10.sim_1.fq -R TAIR10.sim_2.fq -o Atria-consensus --no-tail-n-trim --max-n=-1 --no-quality-trim --no-length-filtration --adapter1 AGATCGGAAGAGCACACGTCTGAACTCCAGTCA --adapter2 AGATCGGAAGAGCGTCGTGTAGGGAAAGAGTGT --threads 1 |
| 1 | AdapterRemoval | 26.35 | 26.35 | 322.19 | 15.82 | 100.0% | 338.05 | 10,572 | AdapterRemoval --file1 TAIR10.sim_1.fq --file2 TAIR10.sim_2.fq --basename AdapterRemoval-3/adapterremoval --adapter1 AGATCGGAAGAGCACACGTCTGAACTCCAGTCA --adapter2 AGATCGGAAGAGCGTCGTGTAGGGAAAGAGTGT --mm 3 --minlength 0 --threads 1 |
| 1 | Skewer | 13.44 | 13.41 | 639.41 | 23.27 | 99.8% | 663.98 | 3,848 | skewer --quiet -x AGATCGGAAGAGCACACGTCTGAACTCCAGTCA -y AGATCGGAAGAGCGTCGTGTAGGGAAAGAGTGT -m pe -l 0 -o Skewer/Skewer TAIR10.sim_1.fq TAIR10.sim_2.fq --threads 1 |
| 1 | Trim Galore | 7.04 | 8.32 | 1,184.12 | 80.57 | 118.1% | 1,070.46 | 16,480 | trim_galore --cores 1 --quality 0 -o TrimGalore --adapter AGATCGGAAGAGCACACGTCTGAACTCCAGTCA --adapter2 AGATCGGAAGAGCGTCGTGTAGGGAAAGAGTGT -e 0.1 --stringency 1 --max_n 100 --length 0 --paired TAIR10.sim_1.fq TAIR10.sim_2.fq |
| 1 | Trimmomatic | 33.22 | 33.73 | 249.17 | 18.96 | 101.5% | 264.10 | 1,318,916 | java -jar /usr/software/Trimmomatic-0.39/trimmomatic-0.39.jar PE -threads 1 -phred33 TAIR10.sim_1.fq TAIR10.sim_2.fq Trimmomatic/out-pair1.paired.fq Trimmomatic/out-pair1.unpaired.fq Trimmomatic/out-pair2.paired.fq Trimmomatic/out-pair2.unpaired.fq ILLUMINACLIP:adapters.fa:2:30:10:1:TRUE:keepBothReads MINLEN:1 |
| 1 | Ktrim | 139.85 | 139.48 | 49.81 | 13.88 | 99.7% | 63.86 | 182,240 | ktrim -1 TAIR10.sim_1.fq -2 TAIR10.sim_2.fq -t 1 -p 33 -q 1 -s 10 -a AGATCGGAAGAGCACACGTCTGAACTCCAGTCA -b AGATCGGAAGAGCGTCGTGTAGGGAAAGAGTGT -o Ktrim/ktrim |
| 1 | Fastp | 9.75 | 13.25 | 852.91 | 60.98 | 135.9% | 672.28 | 3,007,664 | fastp --in1 TAIR10.sim_1.fq --in2 TAIR10.sim_2.fq --out1 fastp/out.fastp.r1.fq --out2 fastp/out.fastp.r2.fq -z 6 --adapter_sequence AGATCGGAAGAGCACACGTCTGAACTCCAGTCA --adapter_sequence_r2 AGATCGGAAGAGCGTCGTGTAGGGAAAGAGTGT --disable_trim_poly_g --disable_quality_filtering --disable_length_filtering --thread 1 |
| 1 | Atropos | 4.32 | 4.32 | 2,034.18 | 27.57 | 100.0% | 2,062.24 | 25,676 | atropos trim -a AGATCGGAAGAGCACACGTCTGAACTCCAGTCA -A AGATCGGAAGAGCGTCGTGTAGGGAAAGAGTGT -o Atropos/TAIR10.sim_1.fq.atropos.fq -p Atropos/TAIR10.sim_2.fq.atropos.fq -pe1 TAIR10.sim_1.fq -pe2 TAIR10.sim_2.fq --aligner insert -e 0.1 |
| 2 | Atria | 59.85 | 108.48 | 132.07 | 16.74 | 181.2% | 82.11 | 2,602,296 | atria --no-consensus -r TAIR10.sim_1.fq -R TAIR10.sim_2.fq -o Atria --no-tail-n-trim --max-n=-1 --no-quality-trim --no-length-filtration --adapter1 AGATCGGAAGAGCACACGTCTGAACTCCAGTCA --adapter2 AGATCGGAAGAGCGTCGTGTAGGGAAAGAGTGT --threads 2 |
| 2 | Atria (consensus) | 45.54 | 85.65 | 180.39 | 15.19 | 188.1% | 103.99 | 2,603,552 | atria -r TAIR10.sim_1.fq -R TAIR10.sim_2.fq -o Atria-consensus --no-tail-n-trim --max-n=-1 --no-quality-trim --no-length-filtration --adapter1 AGATCGGAAGAGCACACGTCTGAACTCCAGTCA --adapter2 AGATCGGAAGAGCGTCGTGTAGGGAAAGAGTGT --threads 2 |
| 2 | AdapterRemoval | 25.99 | 51.65 | 325.97 | 16.80 | 198.8% | 172.45 | 21,236 | AdapterRemoval --file1 TAIR10.sim_1.fq --file2 TAIR10.sim_2.fq --basename AdapterRemoval-3/adapterremoval --adapter1 AGATCGGAAGAGCACACGTCTGAACTCCAGTCA --adapter2 AGATCGGAAGAGCGTCGTGTAGGGAAAGAGTGT --mm 3 --minlength 0 --threads 2 |
| 2 | Skewer | 13.09 | 26.01 | 657.88 | 22.64 | 198.8% | 342.39 | 3,952 | skewer --quiet -x AGATCGGAAGAGCACACGTCTGAACTCCAGTCA -y AGATCGGAAGAGCGTCGTGTAGGGAAAGAGTGT -m pe -l 0 -o Skewer/Skewer TAIR10.sim_1.fq TAIR10.sim_2.fq --threads 2 |
| 2 | Trim Galore | 6.66 | 13.87 | 1,225.64 | 112.56 | 208.4% | 642.02 | 46,568 | trim_galore --cores 2 --quality 0 -o TrimGalore --adapter AGATCGGAAGAGCACACGTCTGAACTCCAGTCA --adapter2 AGATCGGAAGAGCGTCGTGTAGGGAAAGAGTGT -e 0.1 --stringency 1 --max_n 100 --length 0 --paired TAIR10.sim_1.fq TAIR10.sim_2.fq |
| 2 | Trimmomatic | 21.22 | 94.76 | 372.36 | 47.38 | 446.5% | 94.00 | 1,322,756 | java -jar /usr/software/Trimmomatic-0.39/trimmomatic-0.39.jar PE -threads 2 -phred33 TAIR10.sim_1.fq TAIR10.sim_2.fq Trimmomatic/out-pair1.paired.fq Trimmomatic/out-pair1.unpaired.fq Trimmomatic/out-pair2.paired.fq Trimmomatic/out-pair2.unpaired.fq ILLUMINACLIP:adapters.fa:2:30:10:1:TRUE:keepBothReads MINLEN:1 |
| 2 | Ktrim | 99.71 | 176.10 | 73.11 | 16.22 | 176.6% | 50.58 | 341,172 | ktrim -1 TAIR10.sim_1.fq -2 TAIR10.sim_2.fq -t 2 -p 33 -q 1 -s 10 -a AGATCGGAAGAGCACACGTCTGAACTCCAGTCA -b AGATCGGAAGAGCGTCGTGTAGGGAAAGAGTGT -o Ktrim/ktrim |
| 2 | Fastp | 9.15 | 23.43 | 898.02 | 75.60 | 256.1% | 380.16 | 3,113,684 | fastp --in1 TAIR10.sim_1.fq --in2 TAIR10.sim_2.fq --out1 fastp/out.fastp.r1.fq --out2 fastp/out.fastp.r2.fq -z 6 --adapter_sequence AGATCGGAAGAGCACACGTCTGAACTCCAGTCA --adapter_sequence_r2 AGATCGGAAGAGCGTCGTGTAGGGAAAGAGTGT --disable_trim_poly_g --disable_quality_filtering --disable_length_filtering --thread 2 |
| 2 | Atropos | 3.13 | 4.28 | 2,690.20 | 151.07 | 136.5% | 2,081.35 | 1,309,164 | atropos trim -a AGATCGGAAGAGCACACGTCTGAACTCCAGTCA -A AGATCGGAAGAGCGTCGTGTAGGGAAAGAGTGT -o Atropos/TAIR10.sim_1.fq.atropos.fq -p Atropos/TAIR10.sim_2.fq.atropos.fq -pe1 TAIR10.sim_1.fq -pe2 TAIR10.sim_2.fq --aligner insert -e 0.1 --threads 2 --preserve-order |
| 4 | Atria | 55.95 | 168.15 | 142.73 | 16.47 | 300.5% | 52.97 | 2,259,312 | atria --no-consensus -r TAIR10.sim_1.fq -R TAIR10.sim_2.fq -o Atria --no-tail-n-trim --max-n=-1 --no-quality-trim --no-length-filtration --adapter1 AGATCGGAAGAGCACACGTCTGAACTCCAGTCA --adapter2 AGATCGGAAGAGCGTCGTGTAGGGAAAGAGTGT --threads 4 |
| 4 | Atria (consensus) | 43.05 | 136.80 | 190.52 | 16.40 | 317.8% | 65.11 | 2,289,432 | atria -r TAIR10.sim_1.fq -R TAIR10.sim_2.fq -o Atria-consensus --no-tail-n-trim --max-n=-1 --no-quality-trim --no-length-filtration --adapter1 AGATCGGAAGAGCACACGTCTGAACTCCAGTCA --adapter2 AGATCGGAAGAGCGTCGTGTAGGGAAAGAGTGT --threads 4 |
| 4 | AdapterRemoval | 23.63 | 86.25 | 358.28 | 18.67 | 365.0% | 103.27 | 44,120 | AdapterRemoval --file1 TAIR10.sim_1.fq --file2 TAIR10.sim_2.fq --basename AdapterRemoval-3/adapterremoval --adapter1 AGATCGGAAGAGCACACGTCTGAACTCCAGTCA --adapter2 AGATCGGAAGAGCGTCGTGTAGGGAAAGAGTGT --mm 3 --minlength 0 --threads 4 |
| 4 | Skewer | 13.09 | 51.65 | 658.65 | 21.92 | 394.7% | 172.44 | 4,240 | skewer --quiet -x AGATCGGAAGAGCACACGTCTGAACTCCAGTCA -y AGATCGGAAGAGCGTCGTGTAGGGAAAGAGTGT -m pe -l 0 -o Skewer/Skewer TAIR10.sim_1.fq TAIR10.sim_2.fq --threads 4 |
| 4 | Trim Galore | 6.78 | 23.06 | 1,206.08 | 108.53 | 340.3% | 386.33 | 41,444 | trim_galore --cores 4 --quality 0 -o TrimGalore --adapter AGATCGGAAGAGCACACGTCTGAACTCCAGTCA --adapter2 AGATCGGAAGAGCGTCGTGTAGGGAAAGAGTGT -e 0.1 --stringency 1 --max_n 100 --length 0 --paired TAIR10.sim_1.fq TAIR10.sim_2.fq |
| 4 | Trimmomatic | 29.81 | 191.63 | 274.28 | 24.55 | 642.9% | 46.48 | 1,341,460 | java -jar /usr/software/Trimmomatic-0.39/trimmomatic-0.39.jar PE -threads 4 -phred33 TAIR10.sim_1.fq TAIR10.sim_2.fq Trimmomatic/out-pair1.paired.fq Trimmomatic/out-pair1.unpaired.fq Trimmomatic/out-pair2.paired.fq Trimmomatic/out-pair2.unpaired.fq ILLUMINACLIP:adapters.fa:2:30:10:1:TRUE:keepBothReads MINLEN:1 |
| 4 | Ktrim | 74.29 | 243.16 | 103.87 | 16.03 | 327.3% | 36.63 | 336,884 | ktrim -1 TAIR10.sim_1.fq -2 TAIR10.sim_2.fq -t 4 -p 33 -q 1 -s 10 -a AGATCGGAAGAGCACACGTCTGAACTCCAGTCA -b AGATCGGAAGAGCGTCGTGTAGGGAAAGAGTGT -o Ktrim/ktrim |
| 4 | Fastp | 9.68 | 43.40 | 840.32 | 79.77 | 448.3% | 205.24 | 3,223,892 | fastp --in1 TAIR10.sim_1.fq --in2 TAIR10.sim_2.fq --out1 fastp/out.fastp.r1.fq --out2 fastp/out.fastp.r2.fq -z 6 --adapter_sequence AGATCGGAAGAGCACACGTCTGAACTCCAGTCA --adapter_sequence_r2 AGATCGGAAGAGCGTCGTGTAGGGAAAGAGTGT --disable_trim_poly_g --disable_quality_filtering --disable_length_filtering --thread 4 |
| 4 | Atropos | 3.01 | 7.66 | 2,797.51 | 157.68 | 254.2% | 1,162.70 | 537,924 | atropos trim -a AGATCGGAAGAGCACACGTCTGAACTCCAGTCA -A AGATCGGAAGAGCGTCGTGTAGGGAAAGAGTGT -o Atropos/TAIR10.sim_1.fq.atropos.fq -p Atropos/TAIR10.sim_2.fq.atropos.fq -pe1 TAIR10.sim_1.fq -pe2 TAIR10.sim_2.fq --aligner insert -e 0.1 --threads 4 --preserve-order |
| 8 | Atria | 50.15 | 216.56 | 160.07 | 17.55 | 431.9% | 41.13 | 2,500,712 | atria --no-consensus -r TAIR10.sim_1.fq -R TAIR10.sim_2.fq -o Atria --no-tail-n-trim --max-n=-1 --no-quality-trim --no-length-filtration --adapter1 AGATCGGAAGAGCACACGTCTGAACTCCAGTCA --adapter2 AGATCGGAAGAGCGTCGTGTAGGGAAAGAGTGT --threads 8 |
| 8 | Atria (consensus) | 39.10 | 179.50 | 210.31 | 17.50 | 459.1% | 49.62 | 2,502,680 | atria -r TAIR10.sim_1.fq -R TAIR10.sim_2.fq -o Atria-consensus --no-tail-n-trim --max-n=-1 --no-quality-trim --no-length-filtration --adapter1 AGATCGGAAGAGCACACGTCTGAACTCCAGTCA --adapter2 AGATCGGAAGAGCGTCGTGTAGGGAAAGAGTGT --threads 8 |
| 8 | AdapterRemoval | 16.09 | 60.28 | 519.04 | 34.51 | 374.6% | 147.76 | 57,540 | AdapterRemoval --file1 TAIR10.sim_1.fq --file2 TAIR10.sim_2.fq --basename AdapterRemoval-3/adapterremoval --adapter1 AGATCGGAAGAGCACACGTCTGAACTCCAGTCA --adapter2 AGATCGGAAGAGCGTCGTGTAGGGAAAGAGTGT --mm 3 --minlength 0 --threads 8 |
| 8 | Skewer | 12.96 | 100.83 | 666.59 | 20.77 | 778.1% | 88.34 | 5,176 | skewer --quiet -x AGATCGGAAGAGCACACGTCTGAACTCCAGTCA -y AGATCGGAAGAGCGTCGTGTAGGGAAAGAGTGT -m pe -l 0 -o Skewer/Skewer TAIR10.sim_1.fq TAIR10.sim_2.fq --threads 8 |
| 8 | Trim Galore | 6.91 | 34.05 | 1,181.85 | 106.31 | 492.4% | 261.61 | 41,688 | trim_galore --cores 8 --quality 0 -o TrimGalore --adapter AGATCGGAAGAGCACACGTCTGAACTCCAGTCA --adapter2 AGATCGGAAGAGCGTCGTGTAGGGAAAGAGTGT -e 0.1 --stringency 1 --max_n 100 --length 0 --paired TAIR10.sim_1.fq TAIR10.sim_2.fq |
| 8 | Trimmomatic | 30.70 | 250.90 | 267.46 | 22.63 | 817.2% | 35.50 | 1,747,588 | java -jar /usr/software/Trimmomatic-0.39/trimmomatic-0.39.jar PE -threads 8 -phred33 TAIR10.sim_1.fq TAIR10.sim_2.fq Trimmomatic/out-pair1.paired.fq Trimmomatic/out-pair1.unpaired.fq Trimmomatic/out-pair2.paired.fq Trimmomatic/out-pair2.unpaired.fq ILLUMINACLIP:adapters.fa:2:30:10:1:TRUE:keepBothReads MINLEN:1 |
| 8 | Ktrim | 40.67 | 226.64 | 200.72 | 18.27 | 557.2% | 39.30 | 334,928 | ktrim -1 TAIR10.sim_1.fq -2 TAIR10.sim_2.fq -t 8 -p 33 -q 1 -s 10 -a AGATCGGAAGAGCACACGTCTGAACTCCAGTCA -b AGATCGGAAGAGCGTCGTGTAGGGAAAGAGTGT -o Ktrim/ktrim |
| 8 | Fastp | 8.68 | 55.55 | 928.18 | 98.04 | 640.1% | 160.33 | 2,911,100 | fastp --in1 TAIR10.sim_1.fq --in2 TAIR10.sim_2.fq --out1 fastp/out.fastp.r1.fq --out2 fastp/out.fastp.r2.fq -z 6 --adapter_sequence AGATCGGAAGAGCACACGTCTGAACTCCAGTCA --adapter_sequence_r2 AGATCGGAAGAGCGTCGTGTAGGGAAAGAGTGT --disable_trim_poly_g --disable_quality_filtering --disable_length_filtering --thread 8 |
| 8 | Atropos | 2.82 | 15.77 | 2,992.63 | 162.34 | 558.4% | 564.96 | 39,300 | atropos trim -a AGATCGGAAGAGCACACGTCTGAACTCCAGTCA -A AGATCGGAAGAGCGTCGTGTAGGGAAAGAGTGT -o Atropos/TAIR10.sim_1.fq.atropos.fq -p Atropos/TAIR10.sim_2.fq.atropos.fq -pe1 TAIR10.sim_1.fq -pe2 TAIR10.sim_2.fq --aligner insert -e 0.1 --threads 8 --preserve-order |
| 16 | Atria | 37.18 | 241.71 | 219.48 | 20.06 | 650.0% | 36.85 | 2,297,348 | atria --no-consensus -r TAIR10.sim_1.fq -R TAIR10.sim_2.fq -o Atria --no-tail-n-trim --max-n=-1 --no-quality-trim --no-length-filtration --adapter1 AGATCGGAAGAGCACACGTCTGAACTCCAGTCA --adapter2 AGATCGGAAGAGCGTCGTGTAGGGAAAGAGTGT --threads 16 |
| 16 | Atria (consensus) | 30.39 | 215.77 | 274.26 | 18.79 | 709.9% | 41.28 | 2,298,936 | atria -r TAIR10.sim_1.fq -R TAIR10.sim_2.fq -o Atria-consensus --no-tail-n-trim --max-n=-1 --no-quality-trim --no-length-filtration --adapter1 AGATCGGAAGAGCACACGTCTGAACTCCAGTCA --adapter2 AGATCGGAAGAGCGTCGTGTAGGGAAAGAGTGT --threads 16 |
| 16 | AdapterRemoval | 12.52 | 47.93 | 655.04 | 56.23 | 382.8% | 185.82 | 72,896 | AdapterRemoval --file1 TAIR10.sim_1.fq --file2 TAIR10.sim_2.fq --basename AdapterRemoval-3/adapterremoval --adapter1 AGATCGGAAGAGCACACGTCTGAACTCCAGTCA --adapter2 AGATCGGAAGAGCGTCGTGTAGGGAAAGAGTGT --mm 3 --minlength 0 --threads 16 |
| 16 | Skewer | 12.56 | 173.63 | 683.48 | 25.94 | 1382.9% | 51.30 | 10,156 | skewer --quiet -x AGATCGGAAGAGCACACGTCTGAACTCCAGTCA -y AGATCGGAAGAGCGTCGTGTAGGGAAAGAGTGT -m pe -l 0 -o Skewer/Skewer TAIR10.sim_1.fq TAIR10.sim_2.fq --threads 16 |
| 16 | Trim Galore | 6.75 | 36.31 | 1,208.08 | 112.13 | 538.2% | 245.28 | 69,480 | trim_galore --cores 16 --quality 0 -o TrimGalore --adapter AGATCGGAAGAGCACACGTCTGAACTCCAGTCA --adapter2 AGATCGGAAGAGCGTCGTGTAGGGAAAGAGTGT -e 0.1 --stringency 1 --max_n 100 --length 0 --paired TAIR10.sim_1.fq TAIR10.sim_2.fq |
| 16 | Trimmomatic | 25.23 | 250.06 | 328.79 | 24.18 | 990.9% | 35.62 | 2,335,444 | java -jar /usr/software/Trimmomatic-0.39/trimmomatic-0.39.jar PE -threads 16 -phred33 TAIR10.sim_1.fq TAIR10.sim_2.fq Trimmomatic/out-pair1.paired.fq Trimmomatic/out-pair1.unpaired.fq Trimmomatic/out-pair2.paired.fq Trimmomatic/out-pair2.unpaired.fq ILLUMINACLIP:adapters.fa:2:30:10:1:TRUE:keepBothReads MINLEN:1 |
| 16 | Ktrim | 21.67 | 217.19 | 391.92 | 19.03 | 1002.1% | 41.01 | 331,812 | ktrim -1 TAIR10.sim_1.fq -2 TAIR10.sim_2.fq -t 16 -p 33 -q 1 -s 10 -a AGATCGGAAGAGCACACGTCTGAACTCCAGTCA -b AGATCGGAAGAGCGTCGTGTAGGGAAAGAGTGT -o Ktrim/ktrim |
| 16 | Fastp | 6.69 | 51.20 | 1,213.88 | 116.80 | 764.8% | 173.98 | 3,038,220 | fastp --in1 TAIR10.sim_1.fq --in2 TAIR10.sim_2.fq --out1 fastp/out.fastp.r1.fq --out2 fastp/out.fastp.r2.fq -z 6 --adapter_sequence AGATCGGAAGAGCACACGTCTGAACTCCAGTCA --adapter_sequence_r2 AGATCGGAAGAGCGTCGTGTAGGGAAAGAGTGT --disable_trim_poly_g --disable_quality_filtering --disable_length_filtering --thread 16 |
| 16 | Atropos | 2.79 | 15.68 | 3,031.15 | 160.49 | 561.7% | 568.17 | 35,120 | atropos trim -a AGATCGGAAGAGCACACGTCTGAACTCCAGTCA -A AGATCGGAAGAGCGTCGTGTAGGGAAAGAGTGT -o Atropos/TAIR10.sim_1.fq.atropos.fq -p Atropos/TAIR10.sim_2.fq.atropos.fq -pe1 TAIR10.sim_1.fq -pe2 TAIR10.sim_2.fq --aligner insert -e 0.1 --threads 16 --preserve-order |
| 32 | Atria | 19.16 | 225.49 | 443.40 | 21.38 | 1176.7% | 39.50 | 2,048,208 | atria --no-consensus -r TAIR10.sim_1.fq -R TAIR10.sim_2.fq -o Atria --no-tail-n-trim --max-n=-1 --no-quality-trim --no-length-filtration --adapter1 AGATCGGAAGAGCACACGTCTGAACTCCAGTCA --adapter2 AGATCGGAAGAGCGTCGTGTAGGGAAAGAGTGT --threads 32 |
| 32 | Atria (consensus) | 16.50 | 228.50 | 520.27 | 19.66 | 1385.1% | 38.98 | 2,054,080 | atria -r TAIR10.sim_1.fq -R TAIR10.sim_2.fq -o Atria-consensus --no-tail-n-trim --max-n=-1 --no-quality-trim --no-length-filtration --adapter1 AGATCGGAAGAGCACACGTCTGAACTCCAGTCA --adapter2 AGATCGGAAGAGCGTCGTGTAGGGAAAGAGTGT --threads 32 |
| 32 | AdapterRemoval | 11.43 | 45.45 | 695.64 | 83.36 | 397.5% | 195.97 | 114,664 | AdapterRemoval --file1 TAIR10.sim_1.fq --file2 TAIR10.sim_2.fq --basename AdapterRemoval-3/adapterremoval --adapter1 AGATCGGAAGAGCACACGTCTGAACTCCAGTCA --adapter2 AGATCGGAAGAGCGTCGTGTAGGGAAAGAGTGT --mm 3 --minlength 0 --threads 32 |
| 32 | Skewer | 9.34 | 171.45 | 921.59 | 32.31 | 1836.2% | 51.95 | 28,696 | skewer --quiet -x AGATCGGAAGAGCACACGTCTGAACTCCAGTCA -y AGATCGGAAGAGCGTCGTGTAGGGAAAGAGTGT -m pe -l 0 -o Skewer/Skewer TAIR10.sim_1.fq TAIR10.sim_2.fq --threads 32 |
| 32 | Trim Galore | 6.09 | 36.49 | 1,358.40 | 104.65 | 599.3% | 244.12 | 127,188 | trim_galore --cores 32 --quality 0 -o TrimGalore --adapter AGATCGGAAGAGCACACGTCTGAACTCCAGTCA --adapter2 AGATCGGAAGAGCGTCGTGTAGGGAAAGAGTGT -e 0.1 --stringency 1 --max_n 100 --length 0 --paired TAIR10.sim_1.fq TAIR10.sim_2.fq |
| 32 | Trimmomatic | 23.94 | 249.57 | 348.00 | 24.08 | 1042.5% | 35.69 | 2,582,988 | java -jar /usr/software/Trimmomatic-0.39/trimmomatic-0.39.jar PE -threads 32 -phred33 TAIR10.sim_1.fq TAIR10.sim_2.fq Trimmomatic/out-pair1.paired.fq Trimmomatic/out-pair1.unpaired.fq Trimmomatic/out-pair2.paired.fq Trimmomatic/out-pair2.unpaired.fq ILLUMINACLIP:adapters.fa:2:30:10:1:TRUE:keepBothReads MINLEN:1 |
| 32 | Ktrim | 10.38 | 203.08 | 836.79 | 21.67 | 1957.3% | 43.86 | 332,276 | ktrim -1 TAIR10.sim_1.fq -2 TAIR10.sim_2.fq -t 32 -p 33 -q 1 -s 10 -a AGATCGGAAGAGCACACGTCTGAACTCCAGTCA -b AGATCGGAAGAGCGTCGTGTAGGGAAAGAGTGT -o Ktrim/ktrim |
| 32 | Atropos | 2.79 | 15.84 | 3,029.19 | 160.27 | 567.2% | 562.36 | 31,300 | atropos trim -a AGATCGGAAGAGCACACGTCTGAACTCCAGTCA -A AGATCGGAAGAGCGTCGTGTAGGGAAAGAGTGT -o Atropos/TAIR10.sim_1.fq.atropos.fq -p Atropos/TAIR10.sim_2.fq.atropos.fq -pe1 TAIR10.sim_1.fq -pe2 TAIR10.sim_2.fq --aligner insert -e 0.1 --threads 32 --preserve-order |

Input and output files are not compressed. Atria (consensus) does both adapter trimming and paired-end consensus call (base correction of overlapped regions). SeqPurge does not support output uncompressed files, so it is not shown in the uncompressed benchmark. Fastp does not support 32 threads, so only 1-16 threads were tested.

**Table S1-2. Trimming speed on the simulated data (gzip compressed data)**

| Threads | Trimmer | Efficiency (M Bases/s/CPU) | Speed (M Bases/s) | User Time (s) | System Time (s) | CPU | Elapsed Time (s) | Max Memory (kB) | Command |
|---|---|---|---|---|---|---|---|---|---|
| 1 | Atria | 3.58 | 3.91 | 2,436.81 | 53.42 | 109.4% | 2,276.45 | 2,421,012 | atria --no-consensus -r TAIR10.sim_1.fq.gz -R TAIR10.sim_2.fq.gz -o Atria --no-tail-n-trim --max-n=-1 --no-quality-trim --no-length-filtration --adapter1 AGATCGGAAGAGCACACGTCTGAACTCCAGTCA --adapter2 AGATCGGAAGAGCGTCGTGTAGGGAAAGAGTGT --threads 1 |
| 1 | Atria (consensus) | 3.52 | 3.84 | 2,476.92 | 54.11 | 109.1% | 2,319.96 | 2,423,976 | atria -r TAIR10.sim_1.fq.gz -R TAIR10.sim_2.fq.gz -o Atria-consensus --no-tail-n-trim --max-n=-1 --no-quality-trim --no-length-filtration --adapter1 AGATCGGAAGAGCACACGTCTGAACTCCAGTCA --adapter2 AGATCGGAAGAGCGTCGTGTAGGGAAAGAGTGT --threads 1 |
| 1 | AdapterRemoval | 3.65 | 3.65 | 2,433.19 | 5.50 | 100.0% | 2,438.82 | 11,576 | AdapterRemoval --file1 TAIR10.sim_1.fq.gz --file2 TAIR10.sim_2.fq.gz --basename AdapterRemoval-3/adapterremoval --adapter1 AGATCGGAAGAGCACACGTCTGAACTCCAGTCA --adapter2 AGATCGGAAGAGCGTCGTGTAGGGAAAGAGTGT --mm 3 --minlength 0 --threads 1 --gzip |
| 1 | Skewer | 2.78 | 8.49 | 3,165.30 | 42.27 | 305.8% | 1,048.92 | 3,664 | skewer --quiet -x AGATCGGAAGAGCACACGTCTGAACTCCAGTCA -y AGATCGGAAGAGCGTCGTGTAGGGAAAGAGTGT -m pe -l 0 -o Skewer/Skewer TAIR10.sim_1.fq.gz TAIR10.sim_2.fq.gz --threads 1 --compress |
| 1 | Trim Galore | 1.36 | 2.93 | 6,452.45 | 111.48 | 216.1% | 3,038.02 | 16,636 | trim_galore --cores 1 --quality 0 -o TrimGalore --adapter AGATCGGAAGAGCACACGTCTGAACTCCAGTCA --adapter2 AGATCGGAAGAGCGTCGTGTAGGGAAAGAGTGT -e 0.1 --stringency 1 --max_n 100 --length 0 --paired TAIR10.sim_1.fq.gz TAIR10.sim_2.fq.gz |
| 1 | Trimmomatic | 3.61 | 3.61 | 2,429.06 | 40.13 | 100.1% | 2,466.01 | 1,304,372 | java -jar /usr/software/Trimmomatic-0.39/trimmomatic-0.39.jar PE -threads 1 -phred33 TAIR10.sim_1.fq.gz TAIR10.sim_2.fq.gz Trimmomatic/out-pair1.paired.fq.gz Trimmomatic/out-pair1.unpaired.fq.gz Trimmomatic/out-pair2.paired.fq.gz Trimmomatic/out-pair2.unpaired.fq.gz ILLUMINACLIP:adapters.fa:2:30:10:1:TRUE:keepBothReads MINLEN:1 |
| 1 | SeqPurge | 2.69 | 8.06 | 3,279.71 | 33.06 | 299.9% | 1,104.76 | 16,336 | SeqPurge -in1 TAIR10.sim_1.fq.gz -in2 TAIR10.sim_2.fq.gz -out1 SeqPurge/TAIR10.sim_1.fq.gz.seqpurge.fq.gz -out2 SeqPurge/TAIR10.sim_2.fq.gz.seqpurge.fq.gz -a1 AGATCGGAAGAGCACACGTCTGAACTCCAGTCA -a2 AGATCGGAAGAGCGTCGTGTAGGGAAAGAGTGT -mep 0.1 -qcut 0 -min_len 0 -summary SeqPurge/seqpurge.summary -threads 1 |
| 1 | Fastp | 2.96 | 24.49 | 2,893.69 | 116.55 | 827.7% | 363.68 | 16,600 | fastp --in1 TAIR10.sim_1.fq.gz --in2 TAIR10.sim_2.fq.gz --out1 fastp/out.fastp.r1.fq.gz --out2 fastp/out.fastp.r2.fq.gz -z 6 --adapter_sequence AGATCGGAAGAGCACACGTCTGAACTCCAGTCA --adapter_sequence_r2 AGATCGGAAGAGCGTCGTGTAGGGAAAGAGTGT --disable_trim_poly_g --disable_quality_filtering --disable_length_filtering --thread 1 |
| 2 | Atria | 3.46 | 7.91 | 2,515.19 | 56.64 | 228.4% | 1,126.18 | 2,773,240 | atria --no-consensus -r TAIR10.sim_1.fq.gz -R TAIR10.sim_2.fq.gz -o Atria --no-tail-n-trim --max-n=-1 --no-quality-trim --no-length-filtration --adapter1 AGATCGGAAGAGCACACGTCTGAACTCCAGTCA --adapter2 AGATCGGAAGAGCGTCGTGTAGGGAAAGAGTGT --threads 2 |
| 2 | Atria (consensus) | 3.40 | 7.76 | 2,560.00 | 56.31 | 227.9% | 1,148.18 | 2,772,940 | atria -r TAIR10.sim_1.fq.gz -R TAIR10.sim_2.fq.gz -o Atria-consensus --no-tail-n-trim --max-n=-1 --no-quality-trim --no-length-filtration --adapter1 AGATCGGAAGAGCACACGTCTGAACTCCAGTCA --adapter2 AGATCGGAAGAGCGTCGTGTAGGGAAAGAGTGT --threads 2 |
| 2 | AdapterRemoval | 3.65 | 7.30 | 2,431.78 | 7.90 | 199.9% | 1,220.29 | 18,504 | AdapterRemoval --file1 TAIR10.sim_1.fq.gz --file2 TAIR10.sim_2.fq.gz --basename AdapterRemoval-3/adapterremoval --adapter1 AGATCGGAAGAGCACACGTCTGAACTCCAGTCA --adapter2 AGATCGGAAGAGCGTCGTGTAGGGAAAGAGTGT --mm 3 --minlength 0 --threads 2 --gzip |
| 2 | Skewer | 2.34 | 8.49 | 3,742.64 | 66.14 | 363.0% | 1,049.12 | 3,944 | skewer --quiet -x AGATCGGAAGAGCACACGTCTGAACTCCAGTCA -y AGATCGGAAGAGCGTCGTGTAGGGAAAGAGTGT -m pe -l 0 -o Skewer/Skewer TAIR10.sim_1.fq.gz TAIR10.sim_2.fq.gz --threads 2 --compress |
| 2 | Trim Galore | 1.43 | 5.87 | 6,029.34 | 217.94 | 411.4% | 1,518.62 | 45,008 | trim_galore --cores 2 --quality 0 -o TrimGalore --adapter AGATCGGAAGAGCACACGTCTGAACTCCAGTCA --adapter2 AGATCGGAAGAGCGTCGTGTAGGGAAAGAGTGT -e 0.1 --stringency 1 --max_n 100 --length 0 --paired TAIR10.sim_1.fq.gz TAIR10.sim_2.fq.gz |
| 2 | Trimmomatic | 2.89 | 8.32 | 3,023.60 | 61.30 | 288.1% | 1,070.80 | 1,339,960 | java -jar /usr/software/Trimmomatic-0.39/trimmomatic-0.39.jar PE -threads 2 -phred33 TAIR10.sim_1.fq.gz TAIR10.sim_2.fq.gz Trimmomatic/out-pair1.paired.fq.gz Trimmomatic/out-pair1.unpaired.fq.gz Trimmomatic/out-pair2.paired.fq.gz Trimmomatic/out-pair2.unpaired.fq.gz ILLUMINACLIP:adapters.fa:2:30:10:1:TRUE:keepBothReads MINLEN:1 |
| 2 | SeqPurge | 3.98 | 15.89 | 2,202.47 | 37.87 | 399.8% | 560.41 | 16,372 | SeqPurge -in1 TAIR10.sim_1.fq.gz -in2 TAIR10.sim_2.fq.gz -out1 SeqPurge/TAIR10.sim_1.fq.gz.seqpurge.fq.gz -out2 SeqPurge/TAIR10.sim_2.fq.gz.seqpurge.fq.gz -a1 AGATCGGAAGAGCACACGTCTGAACTCCAGTCA -a2 AGATCGGAAGAGCGTCGTGTAGGGAAAGAGTGT -mep 0.1 -qcut 0 -min_len 0 -summary SeqPurge/seqpurge.summary -threads 2 |
| 2 | Fastp | 3.03 | 8.63 | 2,920.61 | 21.05 | 285.1% | 1,031.96 | 3,042,404 | fastp --in1 TAIR10.sim_1.fq.gz --in2 TAIR10.sim_2.fq.gz --out1 fastp/out.fastp.r1.fq.gz --out2 fastp/out.fastp.r2.fq.gz -z 6 --adapter_sequence AGATCGGAAGAGCACACGTCTGAACTCCAGTCA --adapter_sequence_r2 AGATCGGAAGAGCGTCGTGTAGGGAAAGAGTGT --disable_trim_poly_g --disable_quality_filtering --disable_length_filtering --thread 2 |
| 4 | Atria | 3.38 | 15.07 | 2,564.93 | 68.38 | 445.7% | 590.86 | 2,300,680 | atria --no-consensus -r TAIR10.sim_1.fq.gz -R TAIR10.sim_2.fq.gz -o Atria --no-tail-n-trim --max-n=-1 --no-quality-trim --no-length-filtration --adapter1 AGATCGGAAGAGCACACGTCTGAACTCCAGTCA --adapter2 AGATCGGAAGAGCGTCGTGTAGGGAAAGAGTGT --threads 4 |
| 4 | Atria (consensus) | 3.32 | 14.79 | 2,609.96 | 68.99 | 444.8% | 602.31 | 2,299,232 | atria -r TAIR10.sim_1.fq.gz -R TAIR10.sim_2.fq.gz -o Atria-consensus --no-tail-n-trim --max-n=-1 --no-quality-trim --no-length-filtration --adapter1 AGATCGGAAGAGCACACGTCTGAACTCCAGTCA --adapter2 AGATCGGAAGAGCGTCGTGTAGGGAAAGAGTGT --threads 4 |
| 4 | AdapterRemoval | 3.37 | 7.40 | 2,639.21 | 7.68 | 219.8% | 1,204.19 | 34,076 | AdapterRemoval --file1 TAIR10.sim_1.fq.gz --file2 TAIR10.sim_2.fq.gz --basename AdapterRemoval-3/adapterremoval --adapter1 AGATCGGAAGAGCACACGTCTGAACTCCAGTCA --adapter2 AGATCGGAAGAGCGTCGTGTAGGGAAAGAGTGT --mm 3 --minlength 0 --threads 4 --gzip |
| 4 | Skewer | 2.12 | 8.48 | 4,070.75 | 128.05 | 399.8% | 1,050.16 | 4,032 | skewer --quiet -x AGATCGGAAGAGCACACGTCTGAACTCCAGTCA -y AGATCGGAAGAGCGTCGTGTAGGGAAAGAGTGT -m pe -l 0 -o Skewer/Skewer TAIR10.sim_1.fq.gz TAIR10.sim_2.fq.gz --threads 4 --compress |
| 4 | Trim Galore | 1.46 | 11.70 | 5,893.69 | 200.33 | 800.7% | 761.05 | 44,352 | trim_galore --cores 4 --quality 0 -o TrimGalore --adapter AGATCGGAAGAGCACACGTCTGAACTCCAGTCA --adapter2 AGATCGGAAGAGCGTCGTGTAGGGAAAGAGTGT -e 0.1 --stringency 1 --max_n 100 --length 0 --paired TAIR10.sim_1.fq.gz TAIR10.sim_2.fq.gz |
| 4 | Trimmomatic | 2.92 | 8.35 | 2,994.21 | 59.96 | 286.2% | 1,067.08 | 1,334,864 | java -jar /usr/software/Trimmomatic-0.39/trimmomatic-0.39.jar PE -threads 4 -phred33 TAIR10.sim_1.fq.gz TAIR10.sim_2.fq.gz Trimmomatic/out-pair1.paired.fq.gz Trimmomatic/out-pair1.unpaired.fq.gz Trimmomatic/out-pair2.paired.fq.gz Trimmomatic/out-pair2.unpaired.fq.gz ILLUMINACLIP:adapters.fa:2:30:10:1:TRUE:keepBothReads MINLEN:1 |
| 4 | SeqPurge | 4.73 | 24.49 | 1,830.09 | 52.50 | 517.6% | 363.70 | 16,384 | SeqPurge -in1 TAIR10.sim_1.fq.gz -in2 TAIR10.sim_2.fq.gz -out1 SeqPurge/TAIR10.sim_1.fq.gz.seqpurge.fq.gz -out2 SeqPurge/TAIR10.sim_2.fq.gz.seqpurge.fq.gz -a1 AGATCGGAAGAGCACACGTCTGAACTCCAGTCA -a2 AGATCGGAAGAGCGTCGTGTAGGGAAAGAGTGT -mep 0.1 -qcut 0 -min_len 0 -summary SeqPurge/seqpurge.summary -threads 4 |
| 4 | Fastp | 2.99 | 8.62 | 2,940.37 | 35.08 | 288.1% | 1,032.92 | 3,161,256 | fastp --in1 TAIR10.sim_1.fq.gz --in2 TAIR10.sim_2.fq.gz --out1 fastp/out.fastp.r1.fq.gz --out2 fastp/out.fastp.r2.fq.gz -z 6 --adapter_sequence AGATCGGAAGAGCACACGTCTGAACTCCAGTCA --adapter_sequence_r2 AGATCGGAAGAGCGTCGTGTAGGGAAAGAGTGT --disable_trim_poly_g --disable_quality_filtering --disable_length_filtering --thread 4 |
| 8 | Atria | 3.31 | 27.24 | 2,625.60 | 67.75 | 823.8% | 326.96 | 2,620,340 | atria --no-consensus -r TAIR10.sim_1.fq.gz -R TAIR10.sim_2.fq.gz -o Atria --no-tail-n-trim --max-n=-1 --no-quality-trim --no-length-filtration --adapter1 AGATCGGAAGAGCACACGTCTGAACTCCAGTCA --adapter2 AGATCGGAAGAGCGTCGTGTAGGGAAAGAGTGT --threads 8 |
| 8 | Atria (consensus) | 3.25 | 26.75 | 2,675.96 | 68.32 | 824.2% | 332.97 | 2,617,968 | atria -r TAIR10.sim_1.fq.gz -R TAIR10.sim_2.fq.gz -o Atria-consensus --no-tail-n-trim --max-n=-1 --no-quality-trim --no-length-filtration --adapter1 AGATCGGAAGAGCACACGTCTGAACTCCAGTCA --adapter2 AGATCGGAAGAGCGTCGTGTAGGGAAAGAGTGT --threads 8 |
| 8 | Trim Galore | 1.49 | 22.66 | 5,814.96 | 155.89 | 1518.8% | 393.13 | 48,020 | trim_galore --cores 8 --quality 0 -o TrimGalore --adapter AGATCGGAAGAGCACACGTCTGAACTCCAGTCA --adapter2 AGATCGGAAGAGCGTCGTGTAGGGAAAGAGTGT -e 0.1 --stringency 1 --max_n 100 --length 0 --paired TAIR10.sim_1.fq.gz TAIR10.sim_2.fq.gz |
| 8 | SeqPurge | 2.92 | 18.56 | 2,454.65 | 597.81 | 636.2% | 479.83 | 16,252 | SeqPurge -in1 TAIR10.sim_1.fq.gz -in2 TAIR10.sim_2.fq.gz -out1 SeqPurge/TAIR10.sim_1.fq.gz.seqpurge.fq.gz -out2 SeqPurge/TAIR10.sim_2.fq.gz.seqpurge.fq.gz -a1 AGATCGGAAGAGCACACGTCTGAACTCCAGTCA -a2 AGATCGGAAGAGCGTCGTGTAGGGAAAGAGTGT -mep 0.1 -qcut 0 -min_len 0 -summary SeqPurge/seqpurge.summary -threads 8 |
| 16 | Atria | 3.23 | 45.68 | 2,693.23 | 64.82 | 1414.5% | 194.98 | 2,541,276 | atria --no-consensus -r TAIR10.sim_1.fq.gz -R TAIR10.sim_2.fq.gz -o Atria --no-tail-n-trim --max-n=-1 --no-quality-trim --no-length-filtration --adapter1 AGATCGGAAGAGCACACGTCTGAACTCCAGTCA --adapter2 AGATCGGAAGAGCGTCGTGTAGGGAAAGAGTGT --threads 16 |
| 16 | Atria (consensus) | 3.17 | 44.97 | 2,744.78 | 63.21 | 1417.8% | 198.05 | 2,541,872 | atria -r TAIR10.sim_1.fq.gz -R TAIR10.sim_2.fq.gz -o Atria-consensus --no-tail-n-trim --max-n=-1 --no-quality-trim --no-length-filtration --adapter1 AGATCGGAAGAGCACACGTCTGAACTCCAGTCA --adapter2 AGATCGGAAGAGCGTCGTGTAGGGAAAGAGTGT --threads 16 |
| 16 | Trim Galore | 1.27 | 26.18 | 6,894.45 | 139.49 | 2067.6% | 340.20 | 40,784 | trim_galore --cores 16 --quality 0 -o TrimGalore --adapter AGATCGGAAGAGCACACGTCTGAACTCCAGTCA --adapter2 AGATCGGAAGAGCGTCGTGTAGGGAAAGAGTGT -e 0.1 --stringency 1 --max_n 100 --length 0 --paired TAIR10.sim_1.fq.gz TAIR10.sim_2.fq.gz |
| 16 | SeqPurge | 1.86 | 12.92 | 3,968.28 | 815.93 | 694.2% | 689.16 | 17,072 | SeqPurge -in1 TAIR10.sim_1.fq.gz -in2 TAIR10.sim_2.fq.gz -out1 SeqPurge/TAIR10.sim_1.fq.gz.seqpurge.fq.gz -out2 SeqPurge/TAIR10.sim_2.fq.gz.seqpurge.fq.gz -a1 AGATCGGAAGAGCACACGTCTGAACTCCAGTCA -a2 AGATCGGAAGAGCGTCGTGTAGGGAAAGAGTGT -mep 0.1 -qcut 0 -min_len 0 -summary SeqPurge/seqpurge.summary -threads 16 |
| 32 | Atria | 3.01 | 58.09 | 2,899.15 | 62.37 | 1931.5% | 153.33 | 2,528,128 | atria --no-consensus -r TAIR10.sim_1.fq.gz -R TAIR10.sim_2.fq.gz -o Atria --no-tail-n-trim --max-n=-1 --no-quality-trim --no-length-filtration --adapter1 AGATCGGAAGAGCACACGTCTGAACTCCAGTCA --adapter2 AGATCGGAAGAGCGTCGTGTAGGGAAAGAGTGT --threads 32 |
| 32 | Atria (consensus) | 2.95 | 57.41 | 2,951.99 | 62.65 | 1943.2% | 155.14 | 2,528,608 | atria -r TAIR10.sim_1.fq.gz -R TAIR10.sim_2.fq.gz -o Atria-consensus --no-tail-n-trim --max-n=-1 --no-quality-trim --no-length-filtration --adapter1 AGATCGGAAGAGCACACGTCTGAACTCCAGTCA --adapter2 AGATCGGAAGAGCGTCGTGTAGGGAAAGAGTGT --threads 32 |
| 32 | Trim Galore | 1.25 | 25.68 | 6,981.19 | 168.58 | 2061.8% | 346.78 | 38,356 | trim_galore --cores 32 --quality 0 -o TrimGalore --adapter AGATCGGAAGAGCACACGTCTGAACTCCAGTCA --adapter2 AGATCGGAAGAGCGTCGTGTAGGGAAAGAGTGT -e 0.1 --stringency 1 --max_n 100 --length 0 --paired TAIR10.sim_1.fq.gz TAIR10.sim_2.fq.gz |
| 32 | SeqPurge | 1.98 | 4.94 | 4,330.20 | 165.56 | 249.5% | 1,802.15 | 537,256 | SeqPurge -in1 TAIR10.sim_1.fq.gz -in2 TAIR10.sim_2.fq.gz -out1 SeqPurge/TAIR10.sim_1.fq.gz.seqpurge.fq.gz -out2 SeqPurge/TAIR10.sim_2.fq.gz.seqpurge.fq.gz -a1 AGATCGGAAGAGCACACGTCTGAACTCCAGTCA -a2 AGATCGGAAGAGCGTCGTGTAGGGAAAGAGTGT -mep 0.1 -qcut 0 -min_len 0 -summary SeqPurge/seqpurge.summary -threads 32 |

Input and output files are compressed. Atria (consensus) does both adapter trimming and paired-end consensus call (base correction of overlapped regions). In the trimming for compressed data, the speed of AdapterRemoval, Skewer, Fastp, and Trimmomatic kept constant when the number of threads increased from 4 to 32, so we only benchmarked those trimmers using 1, 2, and 4 threads. Atropos was too slow to trim compressed data, and Ktrim did not support output compressed files, so they are not shown in the compressed benchmark.
